# A Singular Base Editing Platform for Polyfunctional Multiplex Engineering of Immune Cells

**DOI:** 10.1101/2025.07.11.664404

**Authors:** Joseph G. Skeate, Nicholas J. Slipek, Walker S. Lahr, Shambojit Roy, Bryce J. Wick, Erin M. Stelljes, Alexandria K. Gilkey, Prateek P. Thenge, Miechaleen D. Diers, Bibekananda Kar, Joshua B. Krueger, Ethan M. Niemeyer, Cara-lin Lonetree, Mitchell G. Kluesner, Jason B Bell, Kendell Clement, Paolo Provenzano, Branden S. Moriarity, Beau R. Webber

## Abstract

Current methods to engineer antigen-specific receptors rely on randomly integrating vectors or double-strand break induced targeted integration, both of which pose safety risks. To implement an all-in-one tool for multiplex knockout (KO) and knock in (KI), we expand the use of cytosine and adenine base editor (ABE) nickase activity to stimulate homology-directed repair (HDR) and insert clinically relevant chimeric antigen receptors (CARs) into specific loci. Through a novel sgRNA design strategy and a recombinant adeno-associated virus (rAAV) delivered DNA template, we enhanced the efficiency of ABE8e-stimulated HDR in human T cells. By combining KI of CD19-, CD33-, or mesothelin-targeting CARs with >95% quadplex gene KO (*B2M/CD3ε/PDCD1/CISH*), we achieve single-step generation of highly functional off-the-shelf CAR T cell products with enhanced function. Importantly, we found no detectable translocations or significant off-target edits and demonstrated efficacy against multiple cancer lines, and a suppressive 3D spheroid culture model. This efficient engineering process of Iterative <u>N</u>icking for <u>S</u>ynchronous <u>E</u>ngineered <u>R</u>eprogramming of <u>T</u> cells (INSERT) establishes a safe, simplified platform for advanced therapeutic CAR T engineering.

## Introduction

As we uncover more of the immune landscape that impacts the functionality of cellular therapies for cancer, increasingly complex engineering strategies for chimeric antigen receptor (CAR)- and T cell receptor (TCR)-based therapies are being sought to improve performance and realize the promise of an off-the-shelf treatment option through genome editing approaches (reviewed in^1^). While engineering methods have been refined to enhance the efficiency of simultaneous gene edits using TALENs and CRISPR/Cas9, the introduction of multiple double-strand breaks (DSBs) poses significant risks, including genotoxicity and chromosomal translocations that compromise the safety and function of cellular products^2–8^. Base editing (BE) offers an unprecedented ability to enable precise multiplex genome modifications without introducing genotoxic DSBs, thus minimizing the risk of unintended chromosomal alterations, including translocations and chromothripsis events^5,9,10^. Recent publications from our laboratory and others have described the broad utility of BE technology for genome editing in various cell types, including primary human immune cells (reviewed in^11^). In recent studies, we demonstrated that targeting splice donor (SD) and splice acceptor (SA) sites could efficiently knockout nearly any protein coding gene^12–14^. We and others used this approach to demonstrate highly efficient, translocation-free multiplex gene editing in primary human T cells and NK cells, highlighting the utility of BE as a safe and effective platform for multiplex genetic modification of primary lymphocytes^15–17^.

For generating and testing next generation cellular therapies it has become common to combine gene knockouts with transgene insertion using transposon-, lenti-, and retroviral-mediated methods (reviewed in^11^). While promising, it remains that these approaches rely on randomly integrating vectors for CAR integration and require numerous molecular components. Recently, there have been documented cases of insertional mutagenesis using these methods, which has prompted the FDA to issue a warning for several commercial CAR T products^18–22^. Though the incidence of secondary malignancies arising from a T-cell origin, let alone manufactured CAR T products, remains incredibly low (<0.1%)^18–22^. Given the associated risks of cellular transformation, site-specific engineering methods should be sought to improve safety and functionality of transgene insertion^23–26^. Creatively, Glaser *et al.* used orthogonal Cas systems to create site-specific KI (∼30%) at the T cell receptor alpha constant (*TRAC*) locus using Cas12a with simultaneous KO of beta-2 microglobulin (B2M, ∼90% KO) and class II major histocompatibility complex transactivator (CIITA, ∼90% KO) using a SpCas9-BE^27^. Importantly, they showed that the DSB induction by the Cas12a system did not lead to translocations at any of the other edited sites; however, any approach using one or more DSBs creates a risk for large deletions, translocations, and unintended genetic alterations at the on-target site^9,28^.

It has previously been shown that staggered or juxtaposed single-strand nicks using Cas9 nickase (nCas9) can induce HDR while reducing off-target mutagenesis by 50- to 1000-fold^29,30^. Deploying a strategy that leverages the nickase function of a base editor to induce HDR insertion of a transgene at a target locus while simultaneously carrying out multiplex base editing of other sites represents a novel approach to greatly simplify manufacturing^30,31^; however, this has yet to be empirically tested and optimized. Here, we demonstrate that adenine base editor (ABE8e) can be used as an all-in-one universal system to drive site-specific KI of CAR expressing genetic cargo via iterative DNA nicking while simultaneously performing highly efficient multiplex gene KO in primary human T cells. This Iterative <u>N</u>icking for <u>S</u>ynchronous <u>E</u>ngineered <u>R</u>eprogramming of <u>T</u> cells (INSERT) method results in the robust generation of highly functional CAR T cells with clinically relevant gene KOs that demonstrate functionality against multiple cancer types.

## Results

### Base editor nickase activity mediates homology-directed repair that is enhanced with iterative nicking sgRNAs

Previous studies have demonstrated KI of transgenes through a dual-nicking strategy where juxtaposed single guide RNAs (sgRNAs) targeting a specific locus can induce HDR of DNA cargo^30,31^. We sought to apply this approach using the Cas9 nickase (nCas9) function of base editors by designing a set of sgRNAs targeting the *AAVS1* locus to create juxtaposed nicks on opposite strands. Using an initial screen of multiple generations of base editors combined with rAAV delivery of a DNA donor template, we observed that as the efficiency of target-base editing increased, our ability to effectively insert cargo decreased, with codon optimized BE4 (coBE4) showing the lowest level of integration (**Figure 1A**). During this initial screening, a high-fidelity adenine base editor, ABE8e, was published^32^. We found that ABE8e stimulated higher efficiency of HDR induction than previous generation BEs. Inspired by the work of others on ‘double-tap’^33^ and recursive editing methods^34,35^, we further enhance KI rates through the introduction of a second set of ‘re-targeting’ sgRNAs complementary to the predicted base edited protospacer sequence for sustained iterative nickase activity (**Figure 1B**). We then carried out a dose-escalation of rAAV to deliver a CD19 CAR-T2A-RQR8 HDR template post-electroporation of ABE8e mRNA alongside 4 sgRNAs targeting the *AAVS1* locus (two targeting and two re-targeting) and observed high frequency KI of the multicistronic transgene cassette (**Figure 1C**). To assess indel occurrence compared to previously published nCas9 methods, we compared ABE8e and nCas9 engineering and surprisingly found that sgRNA sets introducing juxtaposed nicks resulted in a significantly lower level of indels when using ABE8e compared to nCas9 (**Figure 1D**). These data demonstrate that base editors, particularly ABE8e, can be used to mediate HDR using a modified dual-nicking strategy that is enhanced with inclusion of iterative nicking sgRNAs.

**Figure 1.**
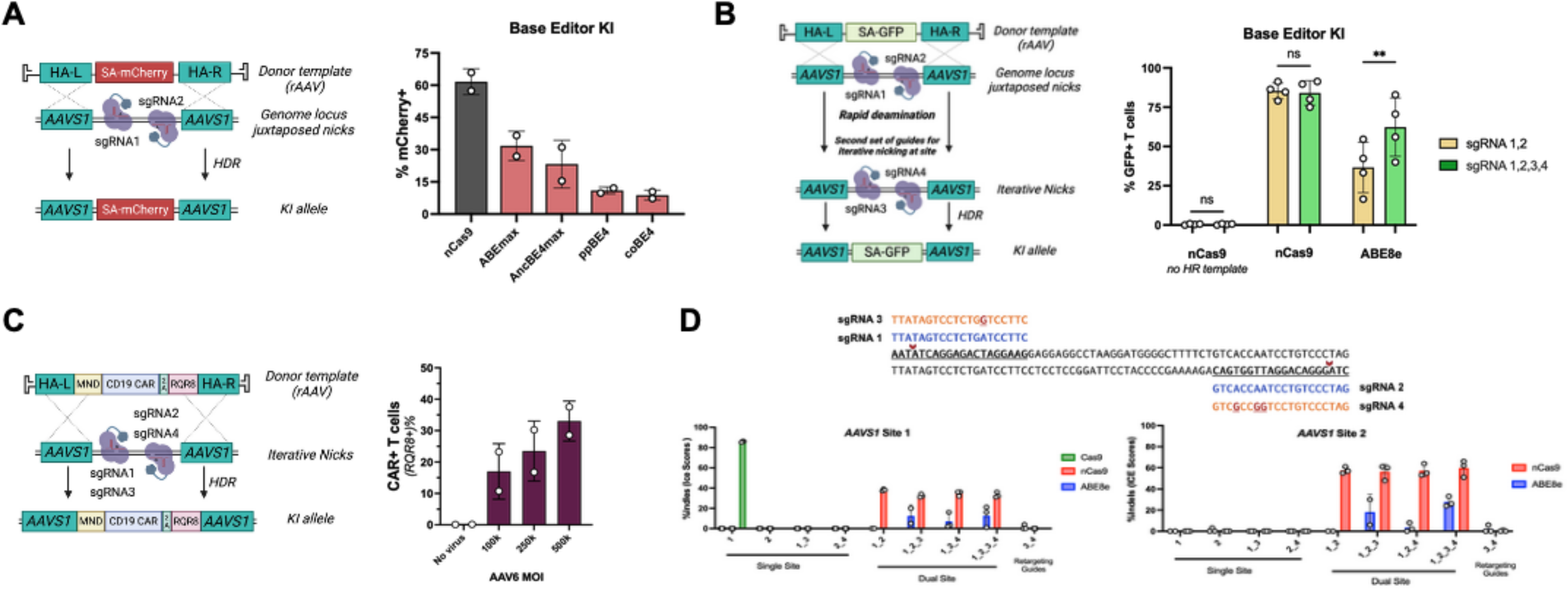
Base editors can be used to facilitate gene KI through homology-directed repair with reduced INDELs at sgRNA target sites. (A, left) Diagram of juxtaposed sgRNA-targeted base editors to facilitate staggered opposite strand nicks and induce HDR. (A, right) Knock-in efficiencies of mCherry cargo into AAVS1 safe harbor locus using juxtaposed sgRNA-targeting (N=2 donors). (B, left) Diagram of juxtaposed and retargeting sgRNA-targeted base editors to facilitate continuous opposite strand nicks and enhance HDR. (B, right) Green fluorescent reporter KI using retargeting sgRNA strategy shows significant improvement of gene KI when using ABE8e (N=4 donors). (C) Successful CD19 CAR RQR8 construct KI at AAVS-1 and expression can be achieved using retargeting sgRNA strategy. (D) Sanger results showing fewer indels are seen at sgRNA target sites with ABE8e compared to nCas9 alone (N=3 donors). *ns = not significant, *p<0.05, unpaired Student’s t-test*.

### Base editor KI efficiency is maintained with contemporaneous multiplex gene KO

After establishing ABE8e paired with iterative nicking for site-directed KI, we next examined whether simultaneous KO of multiple genes is achievable by targeting splice donor (SD) or splice acceptor (SA) sites, as previously published^12,13^. As a proof-of-concept for our INSERT engineering process, we sought to target translationally relevant *beta-2 microglobulin* (*B2M*), which may be used to prevent host versus graft (HvG), CD3 epsilon (*CD3ε*) to inactivate the endogenous TCR signaling to prevent graft versus host (GvH), the internal immune checkpoint cytokine inducible SH2 containing protein (*CISH*), and the surface checkpoint programmed cell death ligand 1 (*PDCD1*)^12,26,36–40^ (**Figure 2A**). Base editor gRNAs to each target were designed using SpliceR^12^ (http://z.umn.edu/spliceR) **(Supplemental Table 1)**. Using our retargeting sgRNA strategy to target the *AAVS1* locus, we were able to successfully KI either a GFP, CD33 CAR, or Mesothelin CAR expressing transgene using rAAV as the HDR template delivery vector (**Figure 2B**). Importantly, KI rates were not impacted with simultaneous three or four gene KO, regardless of transgene cargo, with all base editing rates >95% by NGS analysis and Sanger sequencing (**Figure 2C, Supplemental Table 1, Supplemental Figure 1)**. Flow cytometry analysis of surface expression verified B2M, CD3ε, and PD-1 KO, and Western blots validated CISH KO **(Supplemental Figures 2, 3, and 4)**. Critically, multiplex edited cells did not show impairments in growth or viability during expansion post-engineering **(Supplementary** Figure 5). These data demonstrate successful deployment of INSERT engineering using ABE8e with rAAV HDR template delivery for high fidelity all-in-one multiplex engineering of CAR T cells.

**Figure 2.**
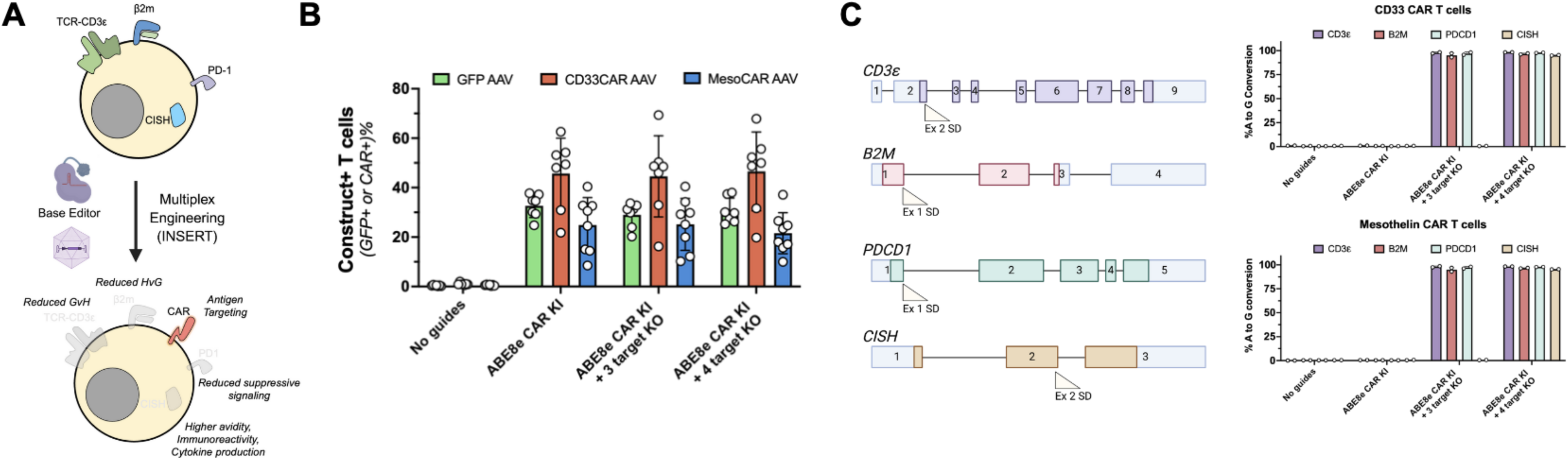
INSERT achieves simultaneous KI of control or CAR cargo with multiplex KO of translational and immunologically relevant genes. (A) Diagram of INSERT engineering outcomes and targeted genes. (B) INSERT-driven KI of control (GFP), CD33 CAR, or Mesothelin CAR is not impacted by the inclusion of targeted gene KO (N=7-8 donors). (C) NGS results showing % A to G conversion of on-target edits for genes targeted for KO from engineered cells.

### Simultaneous KI and KO does not result in detectable translocations or significant off-target editing

Cas9 nuclease-mediated multiplex engineering has shown high-levels of translocations at targeted loci when inducing multiple DSBs simultaneously^10^. Additionally, there is the risk of inducing base editor-installed off-target edits that give rise to single nucleotide variants and ultimately off-target structural variations ^41,42^. It has recently been shown that orthogonal Cas systems can be used to KI cargo while using base-editing to KO multiple genes and this approach does not result in high-levels of translocations, however this method still relies on induction of DSB for knock-in^27^. In contrast, INSERT introduces juxtaposed nicks to induce HDR using the same BE platform that is carrying out the multiplex KO. Previously, we reported that BE-mediated multiplex KO in T cells using cytosine base editors showed near-undetectable levels of translocations when multiplexing targets^15^. To examine whether gene KI simultaneously occurring with multiplex KO results in chromosomal translocations, we developed a ddPCR assay to examine both 5’ and 3’ translocations of all KO genes (*B2M*, *CD3ε, CISH, PDCD1)* with our KI locus (*AAVS1*) (**Figure 3A, Supplementary Table 3)**. Using Cas9 nuclease alongside all 8 sgRNAs we readily detected all predicted translocation events (**Figure 3B).** In contrast, we were unable to detect chromosomal translocations occurring in ABE8e multiplex engineered CD19 CAR T cells, specifically testing for 5’ *AAVS1* or 3’ translocations with target KO genes (**Figure 3B**). In our off-target analysis using rhAmpSeq (Integrated DNA Technologies, Coralville, IA), we analyzed the top ten predicted sites for each guide (IDT CRISPR-Cas9 gRNA checker). We found a single off-target base edit for both *CD3ε* and *PDCD1* sgRNAs (**Figure 4**). For *CD3ε* off-target editing, we found >20% A to G editing at chr9:38085412-38085432, a non-coding region. The *PDCD1* off-target site showed <10% A to G editing within an intronic region (chr15:92446117-92446137) of ST8 alpha-N-acetyl-neuraminide alpha-2,8-sialyltransferase 2 (*ST8SIA2*). These data expand on our previous findings and demonstrate that INSERT engineering results in a CAR T product with near undetectable translocations originating from the site of base editor mediated iterative nicking and very low off-target base editing^15^.

**Figure 3.**
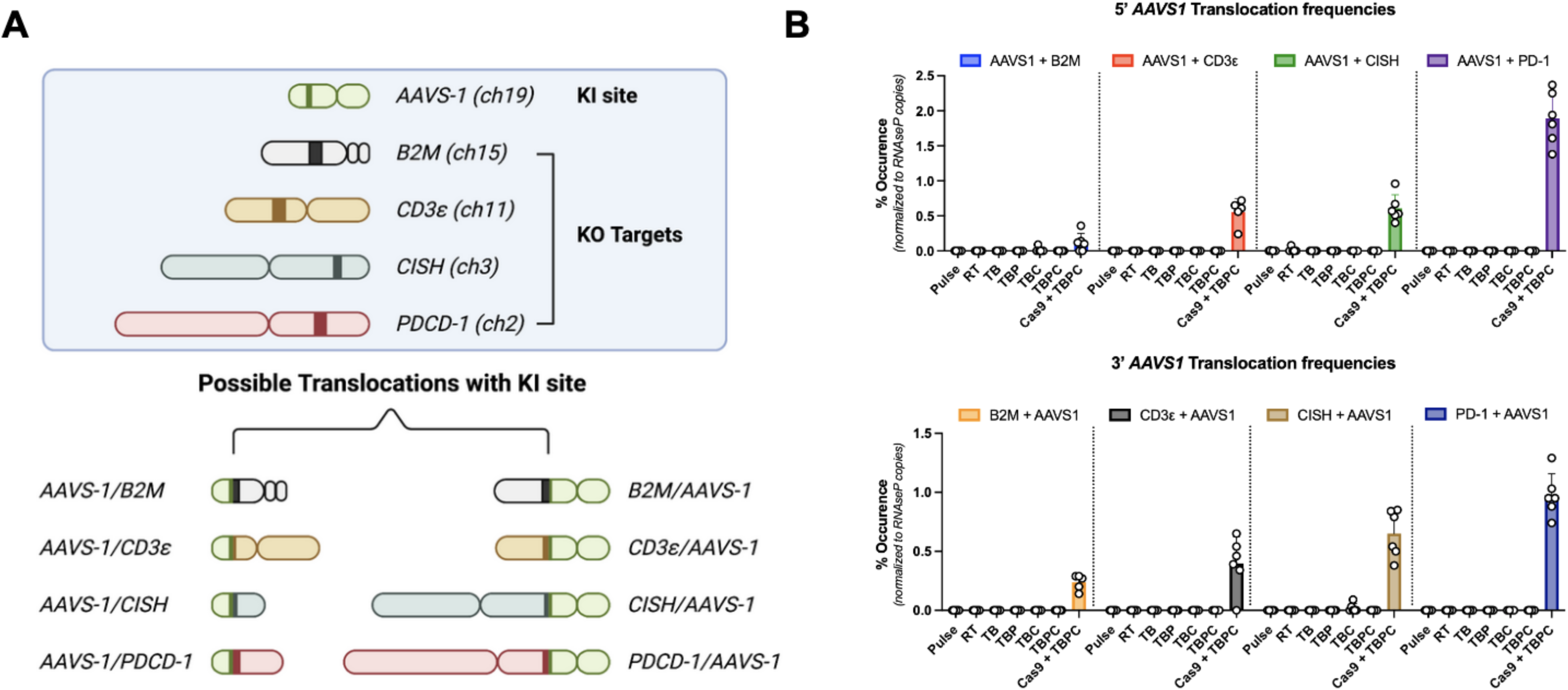
INSERT does not result in detectable translocations. (A) Diagram of possible translocations involving our KI site examined through ddPCR. (B) ddPCR results showing minimal to undetectable translocations in multiplex engineered T cells regardless of the level of edits alongside KI (N=2 donors) (RT = KI only; TB = *CD3ε + B2M* KO; TBP = *CD3ε + B2M + PDCD1* KO; TBC = *CD3ε + B2M + CISH* KO; TBPC = *CD3ε + B2M + CISH + PDCD1* KO).

**Figure 4.**
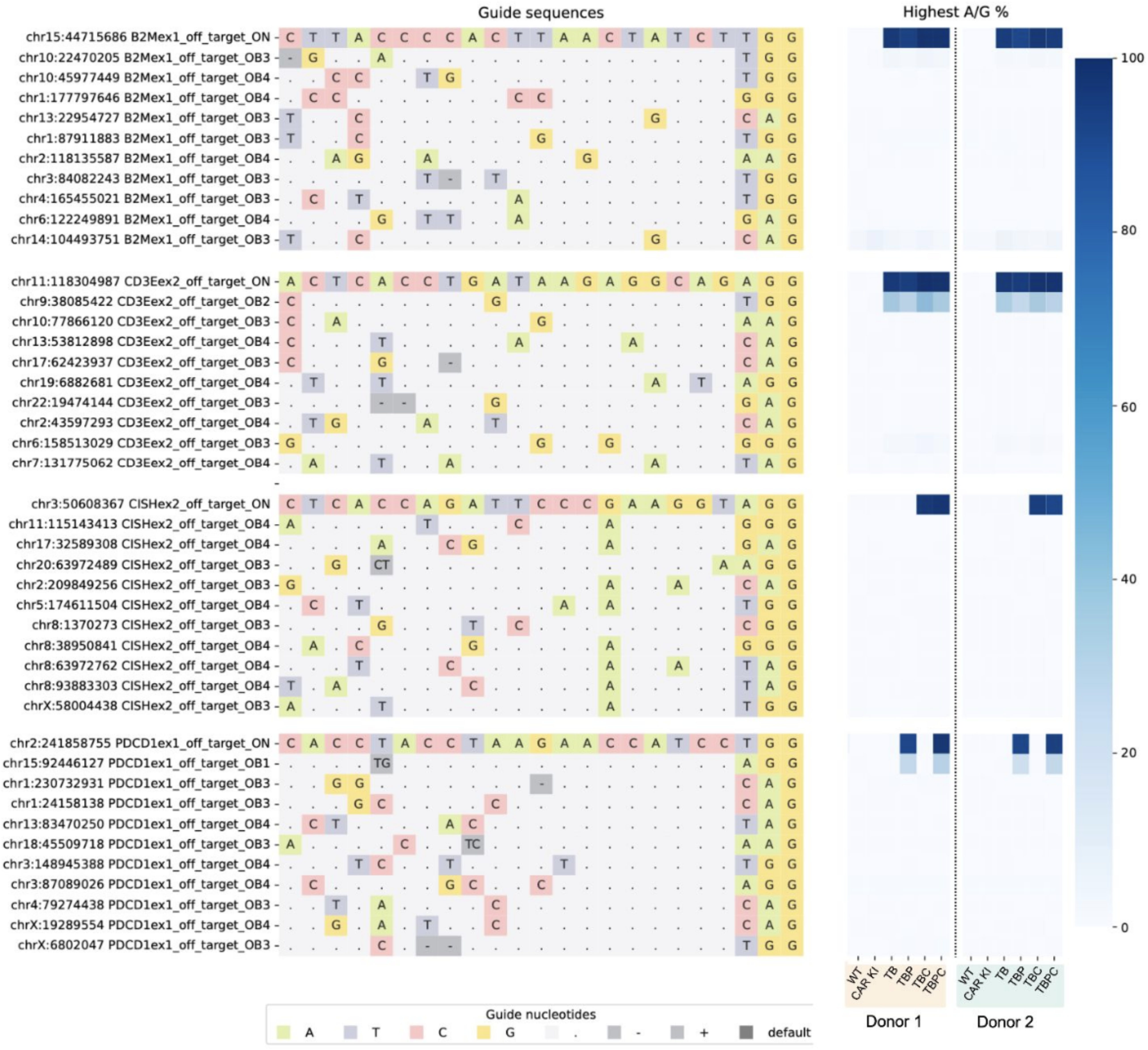
INSERT shows minimal off-target editing. Multiplex engineered cells from two donors were used in rhAmpSeq off-target analysis to identify the top off-target sites susceptible to undesired base editing.

### INSERT CAR T cells show effector function regardless of degree of multiplex KO *in vitro*

We next assessed the cytotoxic function of INSERT engineered CAR T cells and examined whether high-order multiplex KO of genes would lead to negative changes in biology related to INSERT engineering. To this end, CD33 CAR T cells generated through INSERT engineering were assessed for degranulation, cytokine production, and target cell killing using CD33+ MOLM-13 and HL-60 cancer cell lines. Raji cells (CD33-) served as a control for antigen specificity. Additionally, we utilized GFP KI T cells with matching base edits as additional controls to confirm that base edits alone did not impact baseline activity. We found that degranulation (CD107a) and effector cytokine production (IFNγ, TNFɑ) was increased in INSERT CAR T cells compared to controls in co-culture killing assays against CD33+ targets, regardless of the number of genes base edited (T=*CD3ε*, B=*B2M*, P=*PDCD1*, C=*CISH*). No responses were observed in Raji co-cultures or with non-CAR T cells (**Figure 5A, Supplemental Figures 6 & 7)**.

**Figure 5.**
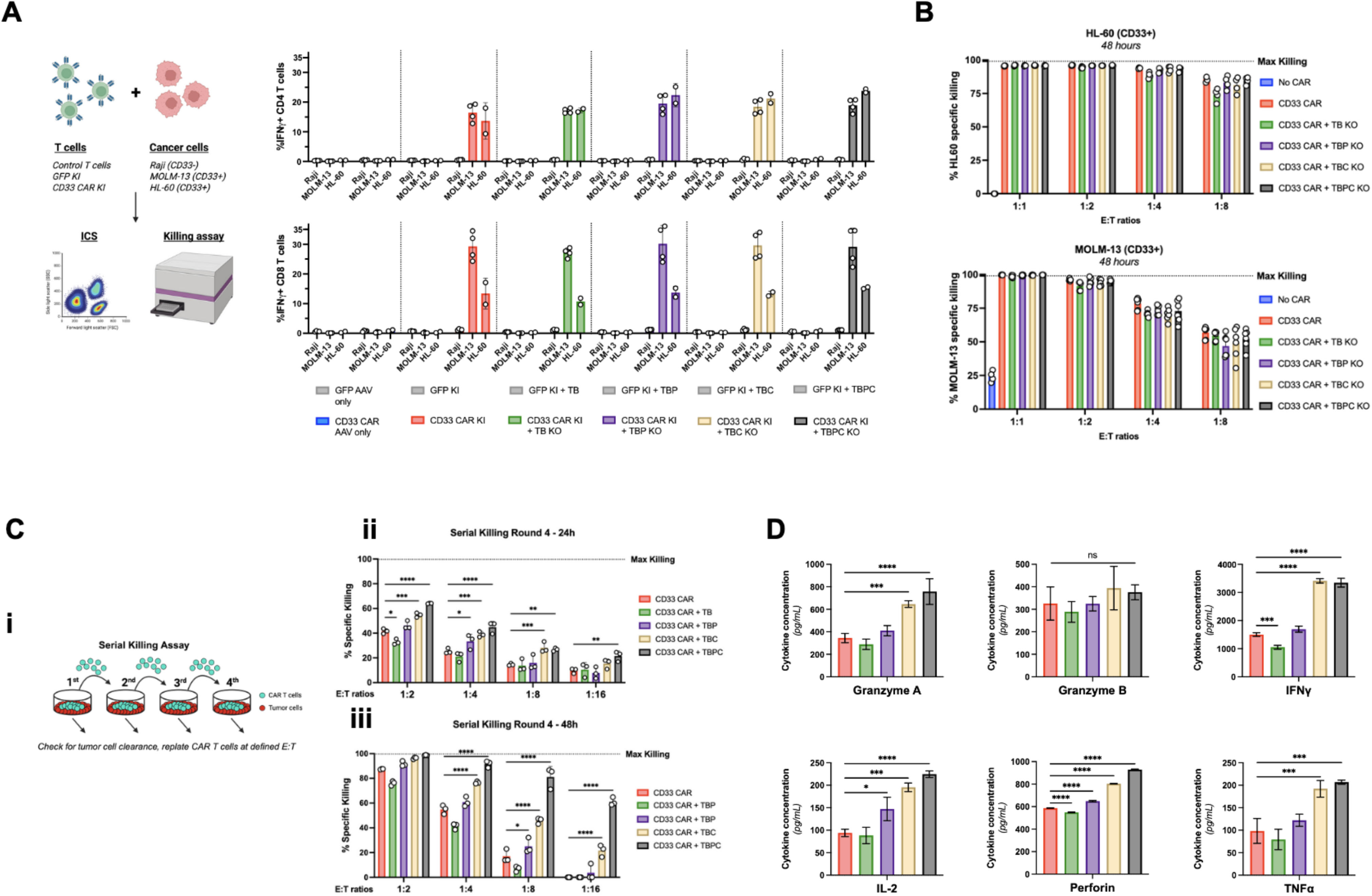
INSERT CD33 CAR T cells show function *in vitro* against tumor lines and demonstrate enhanced function in serial killing when immune checkpoints are KO’d. (A) ICS co-culture assay results showing specific induction of IFNγ against CD33+ cell lines HL-60 and MOLM-13 in CAR T cells but no GFP KI T cells (N=2 donors). (B) CD33 CAR T cells with increasing number of genome edits show function against HL-60 and MOLM-13 at multiple effector to target ratios (E:T) in luciferase-based killing assays. (C) Serial killing assays: i. Diagram of serial killing assay; ii. 24h and iii. 48h readout from 4th round of serial killing of MOLM-13 showing effector function in immune checkpoint multiplex KO’d CAR T cells. (D) Luminex assays showing enhanced cytokine production from supernatants in serial killing assay in immune checkpoint multiplex KO CAR T cells. *ns=not significant, *p<0.05, **p<0.01, ***p<0.001, ****p<0.0001. One-way ANOVA followed by Dunnett’s multiple comparisons test against CAR KI only group (RT)*.

Initial killing assays from 48h co-cultures resulted in robust killing (>95%) of target cells, but did not show differences in effector function between the combination of genome edits at multiple effector to target ratios (E:T) or any response against CD33-negative Raji cells (**Figure 5B, Supplemental Figure 8)**. Next, in an effort to stress-test functional responses, we carried out serial killing through co-culture assays against MOLM-13 and HL-60, adjusting for CAR+ T and cancer cell E:T ratios at the beginning of each round (**Figure 5C**). In the serial killing assays we observed that *B2M*, *CD3ε*, *CISH* or *CISH* + *PDCD1* (TBPC) CD33 CAR T had significantly improved killing of MOLM-13 compared to CAR KI only (**Figure 5C**). Effector cytokine production analyzed via Luminex from the 4th round of serial killing supernatants also showed significant increases in the levels of Granzyme A, IFNγ, IL2, Perforin, and TNFɑ in the multiplex KI/KO groups with *B2M*, *CD3ε, and CISH (TBC)* KO and *TBPC* compared to CAR KI alone (**Figure 5D**). Similar results were seen in serial killing against HL-60 with CAR KI only vs TBPC CAR T **(Supplemental Figure 9)**, and a trend could be seen in OVCAR8 killing with INSERT MesoCAR T cells with increasing gene KO **(Supplemental Figure 10)**. Finally, to examine potential metabolic fitness changes in the different levels of multiplex KO of INSERT CAR T cells, we measured the mitochondrial oxygen consumption rate (OCR) of CD33 CAR T after the first round of target killing through the Seahorse XFe96 Analyzer (Agilent Technologies) and found a higher maximal OCR and spare capacity in our TBPC CD33 CAR T **(Supplemental Figure 11)**. These data demonstrate that INSERT CAR T cells have specific cytotoxic function against target lines and higher degrees of multiplex editing (including *CISH* and/or *PDCD1* KO) do not negatively impact effector function.

### INSERT engineered CD19 CAR T cells show *in vitro* a suppressive 3D model of Burkitt’s Lymphoma

Next, we applied INSERT engineering to produce CD19 CAR T cells and validated on target base edits and protein KO (**Supplemental Figure 12**). Cytotoxic function was assessed against both Raji cells and Raji cells engineered to overexpress hPD-L1 (Raji hPD-L1+) in co-culture ICS assays and in a stress-serial killing assay (**Figure 6A**) (**Supplemental Figure 13 & 14**). We found no significant differences in the level of effector or support cytokine intracellular staining (TNFɑ, IL-2, and IFNγ) or surface CD107a between the different degrees of multiplex KO CAR T (**Supplemental Figure 13**). Although it was not a goal of this engineering study, in the serial killing assays we again saw a significant difference in the multiplex KO CAR T groups TBC and TBPC when compared to the TB group at later rounds of killing (**Supplemental Figure 14**), again highlighting the potential for INSERT engineering to be a useful tool for researchers looking to study the impacts of multi-gene KO.

**Figure 6.**
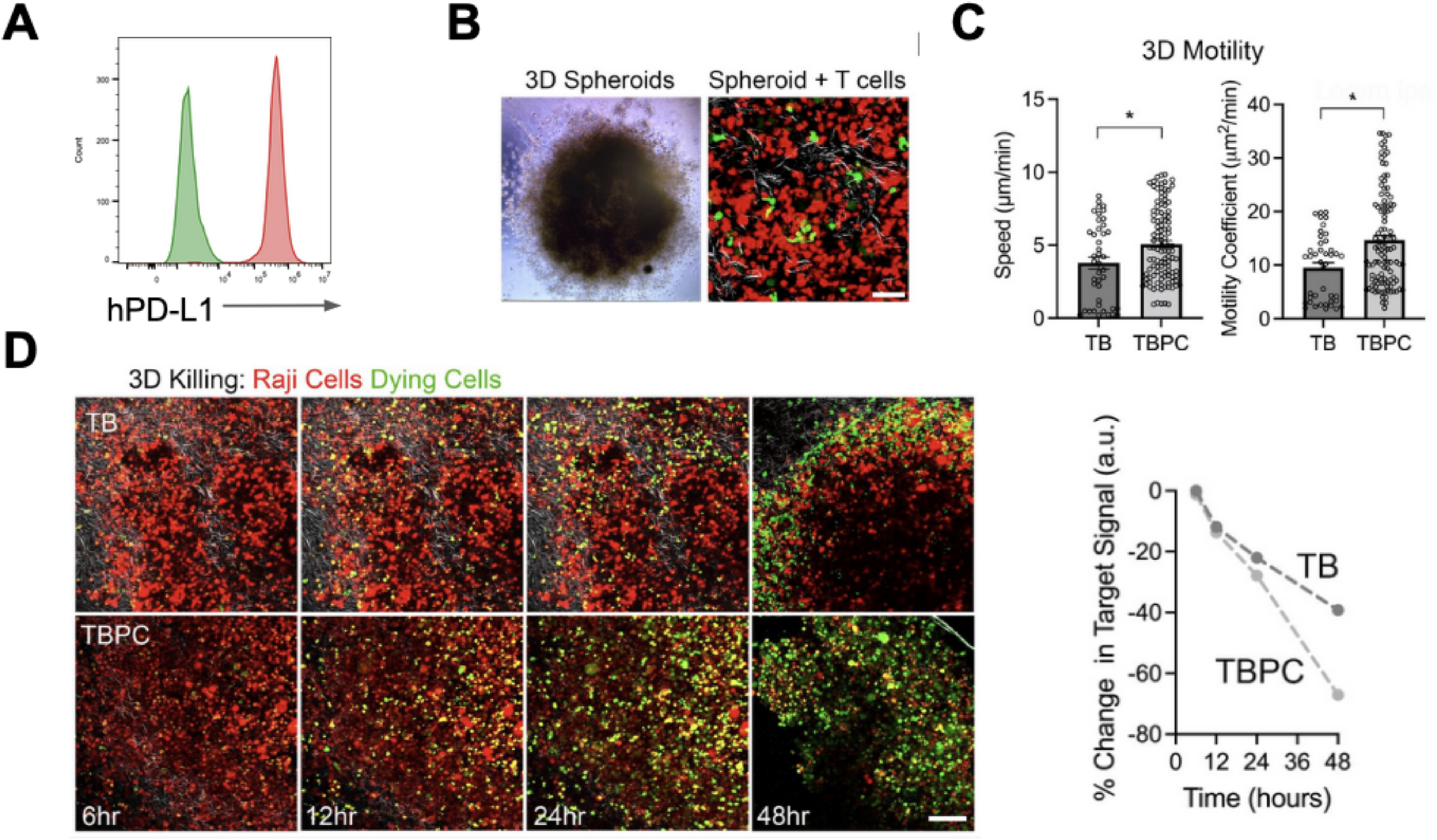
INSERT CD19 CAR T cells show function against an immunosuppressive 3D spheroid model of Burkitt lymphoma. (A) Flow histogram showing Raji cells engineered to overexpress hPD-L1 (**green** = wildtype Raji; **red** = Raji hPD-L1). (B) Example spheroids generated from Raji hPD-L1 cells (left) that are then encased in 3D matrix and therapeutic T cells introduced that migrate to and through the spheroid (**red** = Raji hPD-L1; **green** = T cells; Scale bar = 50 m). (C) Analysis of 3D migration using the persistent random walk model demonstrates that TPBC cells display enhanced speed and overall motility (N=2 donors). (D) 3D killing assay where multiplexed base edited T cells are allowed to engage and kill target cells over time showing increased killing by TBPC T cells (**red** = Raji hPD-L1 cells; **green** = dying cells; Scale bars = 100 µm). *\*p<0.05, unpaired Student’s t-test*.

Finally, in an effort to model how INSERT engineered CAR T would interact with a suppressive lymphoma tumor, we generated Raji-hPD-L1+ spheroids in a 3-dimensional collagen matrix (**Figure 6B**) and assessed the level of infiltration, kinetics of tumor cell death, and CAR T migration through live cell imaging using our minimal (TB) vs our maximal (TBPC) multiplexed INSERT CD19 CAR T. We found that the TBPC CD19 CAR T had significantly increased levels of overall migration in the matrix and increased infiltration into the lymphoma spheroids, complimenting the data seen in the CD33 and CD19 CAR T functional studies (**Figure 6C**). Additionally, kinetic analysis of tumor cell killing clearly demonstrated greater decreases in tumor lymphoid signal in the 48-hour imaging experiments with TBPC CD19 CAR T cells (**Figure 6D**). Overall, these data exemplify that INSERT engineering using ABE8e as an all-in-one modality to facilitate site-specific KI of different CAR cargos and high-fidelity multiplex KO of immunologically related targets generates CAR T cells that are highly functional and may be used to study multigenic modifications in a safe and expedited fashion.

## Discussion

First generation CAR T therapies have seen tremendous success in the clinic against hematologic malignancies (reviewed in^43^); however, therapeutic cell engineering with randomly integrating vectors poses potential risks^44,45^. In late November 2023 a safety communication was released by the Food and Drug Administration which led to the decision to include a warning on all approved BCMA and CD19 CAR T therapies due to the risk of secondary malignancies stemming from the therapy^46^. In addition to safety, costs associated with engineering autologous cell therapies remains high, impacting patient access to these therapies^47,48^. These high costs are not likely to improve as further complexity is incorporated into manufacturing processes, particularly for orthogonal editing strategies where multiple editing enzyme reagents are utilized^27^. For instance, genome engineering strategies using CRISPR/Cas9 systems have demonstrated that it is possible to carry out site-specific transgene KI with high-fidelity using different homology templates for HDR^25,26,49^, but introduction of one or more DSBs with nucleases can result in chromosomal abnormalities, such as translocations or large deletions^50,51^.

As we and others have previously reported, base editors may be employed to generate multiple, highly efficient genome edits with near undetectable indels or translocations in T cells^15,52^. Recently, it’s also been demonstrated that ABE driven multiplex KO CAR T perform superior to Cas9-mediated methods, further driving home the importance of not introducing DSB during the engineering process^17^. Other KI/KO approaches utilized orthogonal Cas systems to mediate KI while also carrying out high-fidelity base editor driven KO at multiple loci^27^. Recently, there has been a novel approach that utilizes customized gRNAs containing MS2 loops^53^ to create a build-your-own aptamer-mediated BE (i.e. Pinpoint) This approach allows nCas9 to carry out KO of target genes as well as juxtaposed nicking to induce HDR-driven knock in at a given locus^54^. While the Pinpoint system achieved effective quadruple gene KO above 50% with successful CAR insertion of ∼20%, it remains that the engineering method requires the co-delivery of nuclease and deaminase mRNA and proprietary sgRNA^54^. In addition to possible cost and GMP scale-up of this custom gRNA architecture, this design has not been as vetted as sgRNAs for off-target or unwanted genome editing impacts.

Here, we demonstrate the use of adenine base editor (ABE8e) and traditional sgRNA architecture for INSERT engineering and achieve >95% on-target gene knockout, up to 70% KI of CAR transgene, no translocations, and incredibly low off-target base editing. We achieved this by employing an HDR-inducing strategy of juxtaposed nicks leveraging ABE8e nickase function and enhanced HDR occurrence through the addition of a second set of sgRNAs that have sequence homology to the base edited target sequence^30,55^. This approach allowed us to KI a variety of cargo through rAAV-mediated HDR template delivery at the *AAVS1* safe-harbor locus, including mCherry, GFP, CD19 CAR, CD33 CAR, and MSLN CAR. Interestingly we found that coBE4, which was used in our previous CAR T engineering study^15^, achieved significantly lower KI than ABE8e and did not show any improvements to KI using the iterative nicking sgRNA approach^15^. Although we did not seek to explore the underlying mechanism behind this observation, it would be a worthwhile follow-up study to examine the editing circuits of each (**Supplemental Figures 15, 16, and 17**) and whether the addition of uracil glycosylase inhibitors to the CBE or the process of base excision repair within the cells prevent iterative nicking required for high efficiency HDR. Identification of the mechanism impeding coBE4 KI may lead to beneficial modifications in the design of future CBEs and CBE-mediated INSERT engineering.

Importantly, we showed that KI does not impair simultaneous multiplex KO of up to 4 targets at greater than 95% efficiency. Surprisingly, at the sgRNA target sites for KI, we observed significantly fewer indels with ABE8e compared with nCas9, suggesting improvements in the safety profile over using nCas9 alone for nickase mediated HDR induction (as in the process utilized by the aforementioned Pinpoint system^54^). The mechanism behind this observation is unclear, but may be due to the reduced frequency of continuous nicking at a target site given the stoichiometric ratio of having 4+ sgRNAs interacting with the base editor versus one or two. Additionally, we were unable to detect chromosomal translocations occurring with the KI site and genes being targeted for KO as well as minimal off-target base editing. As anticipated, we found that regardless of the level of multiplex editing, our engineered cells showed full functionality against cancer cell lines and tumor spheroids. Although not the focus of this study, we show that INSERT engineering enables examination into how targeting multiple genes for KO can impact cell product performance without confounding factors from the engineering process. Specifically, we were able to show improvements to CD33 and CD19 CAR T target killing, motility, cytokine release, and demonstrated the engineered cells performed against 3D spheroids when knocking out multiple immunologically-relevant checkpoints. Given the tumor microenvironment heterogeneity occurring across both cancer types and individual cancers (reviewed in^56^), this platform will enable researchers to rapidly test different combinations of genomic modifications that target negative signaling arising from both intracellular and extracellular factors and tailor cell therapies by the simple addition of additional, off-the-shelf sgRNAs. Together, these observations demonstrate that INSERT engineering enables high-fidelity engineering that next-generation therapies will need to assuage concerns about possible genotoxicities or oncogenic transformation that are being monitored in currently approved CAR T products.

Other improvements to INSERT engineering that would increase speed to clinical translation and bring down overall good manufacturing practice (GMP) engineering costs include use of newly described HDR enhancers, such as 53BP1 inhibiting peptides (IDT) or other HDR enhancing small molecules^57–59^. This could reduce the required MOI of rAAV and allow for the production of more cells per engineering run. Additionally, an investigation into whether INSERT engineering is compatible with novel methods to deliver single stranded DNA templates to facilitate HDR without creating DNA-associated cytotoxicity in engineered cells would allow users to create a fully non-viral, single-step engineering method to generate multiplex edited CAR T cells^49^. If this is achieved, it would allow for a rapidly scalable deployment of testing novel combinations of gene edits for different CAR constructs, which is likely needed to overcome the many therapeutic challenges associated with immunotherapies in solid tumors.

In summary, we report the development of INSERT engineering, an all-in-one system for simultaneous KI and multiplex KO using ABE8e in primary human T cells. This method provides a highly flexible, rapidly deployable platform that allows precise CAR transgene insertion in parallel with multiplex genome engineering to generate therapies against various types of cancers using publicly available reagents. Moving forward, we are actively exploring the application of this platform into the frontier of solid tumors, including enhancement of CAR T cell homing to tumor sites, T cell resistance to oxidative stresses, microenvironment factors, improvements in metabolomics, and more.

## Materials and Methods

### Antibodies

The following antibodies, proteins, and dyes were used: PE- or BV421-conjugated anti-CD33 (clone WM53; BioLegend, San Diego, CA), PE-conjugated anti-mesothelin (clone REA1057; Miltenyi Biotec, Germany), PerCP-conjugated anti-CD19 (clone HIB19; BioLegend), PE- or FitC-conjugated anti-CD3 (clone SK7; BD Biosciences, Franklin Lakes, NJ), BV605-conjugated anti-CD4 (clone RPA-T4; BioLegend), PE-conjugate anti-CD34 to detect RQR8 (clone QBEND/10; Invitrogen, Carlsbad, CA), BV421-conjugated anti-PD-1 (clone EH12.2H7; BioLegend), PE-Cy7-conjugated anti-B2M (clone 2M2, BioLegend) BV650-conjugated anti-CD8 (clone RPA-T8; BioLegend), PE-Cy7-conjugated anti-IL-2 (clone MQ1-17H12; BioLegend), BV421-conjugated anti-IFNγ (clone 4S.B3; BioLegend), APC-conjugated anti-TNFɑ (clone Mab11; BioLegend), BV510-conjugated anti-CD107a (clone H4A3; BD Biosciences), Fixable viability dye eFluor 780 (eBioscience, Carlsbad, CA), Atto 647-conjugated recombinant human CD19 (BioTechne, Minneapolis, MN), Recombinant Human Siglec-3/CD33 Fc Chimera protein (R&D Systems), APC-conjugated anti-human Fc (clone QA10A42; BioLegend), anti-CISH mAb (clone D4D9, Cell Signaling Technologies, Boston, MA), and anti-actin (clone 8H10D10; Cell Signaling Technologies).

### Cell lines

Raji, HL-60, MOLM-13, OVCAR8, and K562 cells were acquired from ATCC. Raji hPD-L1 was generated though lentiviral delivery of a hPD-L1 overexpression vector followed by flow sorting for enrichment of a CD19+, hPD-L1+ population. Routine STR profiling was performed during the study alongside testing for mycoplasma through PCR to monitor for contamination through the Arizona Genomics Core.

### T cell isolation

Bulk-processed peripheral blood T cells were isolated from healthy donor leukopaks with a GMP compliant CliniMACS using TCRα/β GMP MicroBeads (170-070-416, Miltenyi Biotec) and frozen down in Cryostor CS10.

### Cell culture

Primary human T cells were cultured in OpTmizer CTS T cell Expansion SFM containing 2.5% CTS Immune Cell SR (ThermoFisher, Waltham, MA), L-Glutamine (2 mM final concentration), Penicillin/Streptomycin (10 U/mL), N-Acetyl-L-cysteine (10 mM), IL-2 (Peprotech, 300 IU/mL), IL-7 (Peprotech, 5 ng/mL), and IL-15 (Peprotech, 5 ng/mL) at 37 °C and 5% CO2 (Complete OpTimizer). Prior to electroporation, T cells were activated with Dynabeads Human T-Activator CD3/CD28 (ThermoFisher) at a 2:1 bead:cell ratio for 36 hrs. Following electroporations and recovery, T cells were maintained at ∼1 × 10^6^ cells/mL in a 24-well G-Rex (Wilson Wolf, Saint Paul, MN) or 12-well plates during expansions. Target cell lines OVCAR8, K562, MOLM-13, Raji, and Raji-PDL1 cells were cultured in RPMI-1640 supplemented with 10% FBS and Penicillin/Streptomycin (10 U/mL). HL-60 cells were cultured in DMEM + 20% FBS and Penicillin/Streptomycin (10 U/mL).

### All-in-one INSERT CAR T engineering

Thirty-six hours post-activation, dynabeads were magnetically removed and cells were washed once with PBS prior to resuspension in Amaxa primary cell electroporation buffer (P3). 1 × 10^6^ T cells were electroporated with 1 µg of each chemically modified sgRNA (IDT, Coralville, Iowa) and 1.5 µg codon optimized ABE8e-NGG mRNA, codon-optimized *Sp*Cas9 mRNA (TriLink Biotechnologies, San Diego, CA), or CleanCap Cas9 mRNA (TriLink Biotechnologies) with the 4D-nucleofector system (Lonza, Basel, Switzerland) using a 16-well Nucleocuvette kit using the program EO-115. T cells were allowed to recover in electroporation cuvettes for 15 mins at RT before transferring to antibiotic-free medium at 37 °C, 5% CO2 for an additional 30 mins. Cells were then transferred to new 96-well u-bottom plates containing AAV6 harboring our gene KI cargo of interest at MOI indicated.

### Flow phenotyping of engineered T cells

At day 7 or 9 post electroporation, cells were harvested, washed, and incubated with human TruStain FcX and a fixable live/dead dye (eFluor780, 1:500 dilution in PBS) for 15 mins at RT. Afterwards, surface staining for CD3, CD4, CD8, RQR8, PD-1, and B2M was then performed for 20 mins at RT. Cells were washed twice in PBS and analyzed on a CytoFlex S flow cytometer (Beckman Coulter, Brea, CA, USA). Data analysis was performed using FlowJo version 10.10.1 software (FlowJo LLC). For full validation of PD-1 KO in a subset of engineering runs, cells were stimulated with dynabeads and cultured in

### Immunoblot analysis of CISH KO

Proteins were isolated from 1 × 10^6^ cells using complete RIPA buffer with protease and phosphatase inhibitors (Sigma-Aldrich, COEDTAF-RO, P5726, and P0044). Total protein concentration was determined using the Pierce BCA Protein Assay Kit (ThermoFisher, Waltham, MA) according to the manufacturer’s protocol. Cell lysates were adjusted to a concentration of 1 µg/µL and denatured at 95°C for 5 mins. Samples were then analyzed on the JESS platform (ProteinSimple, San Jose, CA) according to the manufacturer’s protocol. Primary antibodies against CISH and actin were used at 1:100 and 1:50 dilutions, respectively, in the kit-supplied buffer. Platform-optimized secondary antibodies were purchased from ProteinSimple.

### Sanger sequencing

Seven days post-electroporation, genomic DNA was isolated from T cells by spin column-based purification according to the manufacturer’s protocol (GeneJet, ThermoFisher). Editing efficiency was accessed by PCR amplification of the targeted loci using AccuPrime Taq DNA Polymerase (Invitrogen, Carlsbad, CA) (**Supplemental Tables 1 and 2**). Sanger sequencing of the PCR amplicons was carried out by Eurofins Genomics. Base editing efficiency was analyzed using EditR (baseeditr.com)^60^, while INDEL frequencies was quantified using Synthego ICE (https://ice.synthego.com/#/).

### NGS

Primers containing Nextera universal primer adapters (Illumina, San Diego, CA) were designed using Primer3Plus to amplify a 375–425 bp region flanking the site of interest. Genomic DNA was PCR-amplified using AccuPrime Taq DNA Polymerase (Invitrogen) with the following thermal cycling condition: [94 °C—2:00]−30 × [94 °C—0:30, 55 °C—0:30, 68 °C—0:30]-[68 °C—5:00]-[4 °C—hold]. Amplicons were extracted from a 1% agarose gel using the QIAquick Gel Extraction Kit (Qiagen, Hilden, Germany). Samples were further amplified with indexing primers and sequenced using a MiSeq 2 × 300 bp run (Illumina, San Diego, CA). A minimum of 20,000 reads were used for INDEL and base editing analysis.

### rhAMPseq off-target analysis

Genomic DNA was purified, and concentration was verified using JetSeq Library Quantification Lo-Rox kit (BIO 68029, Meridian BioScience, Memphis, TN) and Quant-iT PicoGreen (P7589, Invitrogen, Carlsbad, CA). The NGS library was prepared using rhAmpSeq CRISPR Library Kit (10007318, Integrated DNA Technologies) and all samples were treated according to the manufacturer’s protocol. Briefly, 50 ng of genomic DNA was added to the Targeted rhAmp PCR step 1, which contains rhAmpSeq CRISPR forward and Reverse pools. The post-PCR1 product was diluted 1:20 and added to Indexing PCR step 2 using Illumina Nextera XT primers (Illumina, San Diego, CA). Finally, libraries were pooled and purified with AMPure XP beads (Beckman Coulter, Indianapolis, IN). Sequencing was performed on an Illumina MiSeq 2X300 bp paired-end read length (Illumina).

Sequencing reads were analyzed using CRISPRessoPooled^61^. First, reads were aligned to the hg38 genome in genome-only mode using the command: CRISPRessoPooled -x genome/hg38 -r1 fastq.r1.fq -r2 fastq.r2.fq. The genomic sequences of highly covered regions were extracted from the REPORT_READS_ALIGNED_TO_GENOME_ALL_DEPTHS.txt output file and used to assign gRNA on- and off-targets to genomic regions, creating a highly-covered-regions.txt file containing amplicons and their associated gRNA sequence. Next, sequencing reads were re-analyzed with CRISPRessoPooled in genome-and-amplicons mode using the command: CRISPRessoPooled -x genome/hg38 -f highly-covered-regions.txt -r1 fastq.r1.fq -r2 fastq.r2.fq. For each region, CRISPResso outputs included counts of reads containing each nucleotide or indels at every position (Nucleotide_frequency_table.txt and Modification_count_vectors.txt). These data were used to calculate the maximum A>G, C>T, and indel rates at positions 3–17 of the gRNA spacer sequence.

### Intracellular cytokine staining

T cells were thawed and co-cultured with the indicated target cells at a 1:1 effector:target (E:T) ratio in RPMI-1640 supplemented with 10% FBS and Pen/Strep. The cells were incubated at 37 °C with 5% CO2 in the presence of a CD107a antibody for 1 hr followed by a treatment with brefeldin A (10 μg/mL; BD Biosciences) and monensin (0.7 μg/mL; BD Biosciences). After 6 hrs of incubation at 37 °C with 5% CO2, the cells were collected, and incubated with Fixable Viability Dye eFluor780 (eBioscience) and human TruStain FcX (BioLegend) for 10 mins at RT. The cells were then stained with fluorescently labeled antibodies against CD4, CD8, and RQR8 for 20 mins at RT. Afterward, the cells were washed twice in PBS and permeabilized with Fix/Perm (BD Biosciences) for 20 mins at RT. Following permeabilization, the cells were washed with Perm/Wash (BD Biosciences) and stained with fluorescently labeled antibodies against IFNγ, TNFa, and IL-2 for 20 mins at RT. After final washes, the cells were analyzed using a CytoFlex S flow cytometer (Beckman Coulter). Data analysis was performed using FlowJo version 10.10.1 (FlowJo LLC).

### In vitro killing and serial killing assays

Engineered CAR T cells were stress-tested for *in vitro* efficacy against CD33 and CD19-expressing cancer cell lines. Briefly, 50,000 target cells were plated in RPMI-1640 medium supplemented with 10% FBS in 96-well opaque plates. CAR T cells were added at indicated E:T ratios. D-luciferin (D-luc) (Goldbio, St. Louis, MO) was added at indicated time points to assess target cell killing via plate-reader. Specific killing was measured as a percentage of luminescence normalized to the control groups (target-expressing cells alone), with maximum killing defined as the luminescence value of cells treated with 2% Triton-X-100. At time points where target cells were eliminated (indicated by no bioluminescence signal and validated by flow cytometry for target cell removal), CAR T cells from technical replicates were harvested, counted, and phenotyped via flow cytometry for viability (eFluor780), T cell lineage and memory markers (CD45ro, CD62L, CD4, CD8, CD3), CAR expression (RQR8), and activation/exhaustion markers (Ox40, PD-1, TIM-3, LAG-3). A new round of serial killing was then initiated by adding 50,000 target cells to a fresh 96-well U-bottom opaque plate and co-culturing with CAR T cells at indicated E:T ratios.

### Mitochondrial oxygen consumption rate (OCR) analysis

Engineered CAR T collected between the first and second rounds of serial killing against HL-60 cells were analyzed using the Seahorse XFe96 analyzer (Seahorse Bioscience, Billerica, MA, USA). A total of 200,000 cells per well were seeded on poly-D-lysine-coated plates on the same day as the experiment, following the manufacturer’s instructions. The following mitochondrial respiration parameters were measured: (i) basal respiration; (ii) leak respiration, assessed by treating cells with Oligomycin A (Oligo); (iii-iv) maximal respiration, determined by sequential additions of carbonyl cyanide 3-chlorophenylhydrazone (CCCP) (C2759, Sigma) to fully reach mitochondrial reserve capacity; and (v) non-mitochondrial Oxygen consumption, measured after treatment with rotenone and antimycin A.

### Luminex assay for secreted cytokine analysis

Cell culture supernatants were collected at 24 hr timepoints in serial killing assays and clarified by centrifugation at 1,000 × g for 5 mins to remove potential cell debris. Secreted cytokine levels were analyzed using a custom ProcartaPlex human cytokine immunoplex assay (Invitrogen) following the manufacturer’s instructions. Assays were read on a Luminex 200 instrument controlled by BioPlex manager software (version 6.2) collecting a minimum of 50 beads per region of interest.

### Droplet digital polymerase chain reaction translocation analysis

Translocation PCR assays were designed using PrimerQuest software (Integrated DNA Technologies,) with settings optimized for 2 primers plus probe qPCR assay based on custom designed translocation maps. Each sample was run as a duplexed assay consisting of an internal reference primer - probe set targeting RNAseP (HEX) and an experimental primer -probe set (FAM). Primers and probes were ordered from IDT. Reactions were set up using the ddPCR Supermix for Probes (no dUTP) (Bio-Rad, Hercules, CA) with 50 ng of genomic DNA per assay following the manufacturer’s instructions. Droplets were generated and analyzed using a QX200 Droplet-digital PCR system (Bio-Rad, Hercules, CA). Translocation frequency was calculated as the fractional abundance of the experimental target, normalized to the reference sequence (RNAse P, present in two copies per genome) using the QuantaSoft ver 14.0 software (Bio-Rad, Hercules, CA).

### Preparation of bulk collagen gel with spheroids

To prepare bulk collagen gels, the stock collagen (Corning, Corning, NY) was first mixed with 100 mM HEPES buffer (prepared in 2X PBS) on ice. An equal volume of ice cold complete RPMI media (RPMI 1640, 10% FBS, 10 U/mL Pen/Strep) was then added to this mixture to achieve a final volume of 1 mL and a collagen concentration of 3 mg/mL. Raji hPD-L1 spheroids were carefully transferred to 1.5 mL Eppendorf tubes (one spheroid per tube) by gently pipetting out from the wells. The spheroids were then washed twice with 0.5 mL 1X PBS, followed by resuspension in 0.5 mL complete medium. CellTracker Red (Invitrogen) was added to a final concentration of 5 μM to stain spheroids, and tubes were incubated for 10 mins. After incubation, the medium was carefully removed from the tubes, without disturbing the spheroid, which were then washed with PBS twice. Post removal of final wash, tubes were then placed on ice to allow the spheroids to equilibrate to the collagen temperature. 150 µl of cold collagen (3 mg/mL) was then added to each tube, while the remaining 850 µL of collagen was transferred to a 35 mm tissue culture-treated Petri Dish (Thermo Fisher). Dishes were incubated for 12 mins to allow for partial polymerization, Raji hPD-L1 spheroids were then transferred to the partially polymerized collagen gel. The dish was then placed back into the incubator, allowing the collagen to fully polymerize for at least 16 hrs. This method ensured proper embedding of the spheroids within the collagen gel.

### In vitro CAR T cell migration studies with 3D bulk gels

INSERT CD19 CAR T cells were cultured in complete T cell medium (RPMI supplemented with 10% FBS, 10 U/mL Pen/Strep and 30 U/mL IL-2) in round bottom, tissue culture-treated 96-well plates (Corning) at a density of 1 x 10^5^ cells per well. The plates were incubated at 37 °C with 5% CO_2_. On the day of the experiment, T cells were collected and transferred to a 15 mL conical tube. Cell counts and viability were assessed using Trypan blue exclusion using a Thermo Countess II (Thermo Fisher Scientific, Waltham, MA). CellTracker Green (Invitrogen) was added to the T cells to a final concentration of 5 µM and incubated for 10 mins at 37 °C. After incubation, T cells were washed twice with PBS. T cells were suspended in complete L-15 medium (Gibco, 10% FBS, 10 U/mL Pen/Strep). Labeled T cells were then added to completely polymerized collagen bulk gel containing the spheroid at a density of 1 x 10^5^ cells per dish. The dish (spheroid + T cells) was placed inside a 37 °C incubator. After 2 hrs, the bulk gel was washed with PBS to remove any free-floating T cells, and 2 mL of fresh L-15 medium was added to the dish. The dishes containing spheroids and T cells were then imaged for 3 hrs using a multiphoton laser scanning microscope (Prairie Technologies). Imaging was performed at a single NIR wavelength (880 nm) to visualize spheroids (red), TB CAR T cells (green) and collagen (SHG). Images were acquired with a 20X objective, with a frame frequency of 3 mins, with a z-depth of 50 µm, and 2.5 µm per optical section, resulting in 20 sections in the z-direction and 60 time frames. The same procedure was repeated for TBPC CAR T cells.

### In vitro cytotoxicity assay in 3D bulk gels

For cytotoxicity experiments, 1 x 10^5^ unstained INSERT CAR T cells were resuspended in 1 mL of complete L-15 medium and added to the fully polymerized bulk gels containing spheroids. Cytotox Green (Sartorius, Göttingen, Germany), was added to a final concentration of 0.5 µM to label non-viable cells. The dish was then incubated at 37°C in an air incubator for 2 hrs. After incubation, the bulk gel was washed twice with PBS, to remove any free-floating T cells, and 2 mL of fresh L-15 medium was added. Time lapse imaging was performed at 6, 12, 24 and 48 hrs using a multiphoton laser scanning microscope (Prairie Technologies). A NIR laser at 880 nm was used to generate 3 channels for visualization: spheroids (red), dead cancer cells (green), and collagen (SHG). Images were acquired using a 20x objective, with a z-depth of 50 µm, and an optical section thickness of 2.5 µm, resulting in 20 sections per z-stack.

### Image processing and data analysis

Images from the migration experiment were analyzed using ImageJ. Z-sections were compressed using maximum intensity projection across all 60 time frames. T cells were identified and their migration history was tracked across all optical sections and time frames using the TrackMate plugin of Fiji. The 3D speed and motility coefficients of individual T cells were calculated using a persistent random walk model (PRWM) developed by the Provenzano lab^62^, and the mean values for each parameter were reported. T cell cytotoxicity against the Raji spheroids was quantified by calculating the percentage change in fluorescent intensity from the green channel over the time course.

## Acknowledgements

J.G.S. is supported by the T32HL007062-46 Hematology Research Training Program. M.G.K. is supported by Washington Research Foundation via the Achievement Rewards for College Scientists Fellowship. P.P.P acknowledges funding from NIH grants U54CA268069, P01CA254849, R01CA245550, and MBGI-22-009-01-MBG from the American Cancer Society. B.R.W. acknowledges funding from NIH grants R21CA237789, R21AI163731, P01CA254849, P50CA136393, R01AI146009, U54CA268069, Children’s Cancer Research Fund, Cure Childhood Cancer, the Randy Shaver Cancer Research and Community Fund. BSM acknowledges funding from Office of Discovery and Translation, NIH grants (R01AI146009, R01AI161017, P01CA254849, P50CA136393, U24OD026641, U54CA232561, P30CA077598, U54CA268069, DOD grants HT9425-24-1-1005, HT9425-24-1-1002, HT9425-24-1-0231), and Children’s Cancer Research Fund, the Fanconi Anemia Research Fund, and the Randy Shaver Cancer and Community Fund.

