## Supplemental Data for "A Singular Base Editing Platform for Polyfunctional Multiplex Engineering of Immune Cells"

### Supplemental 1: Sanger analysis of on-target edits

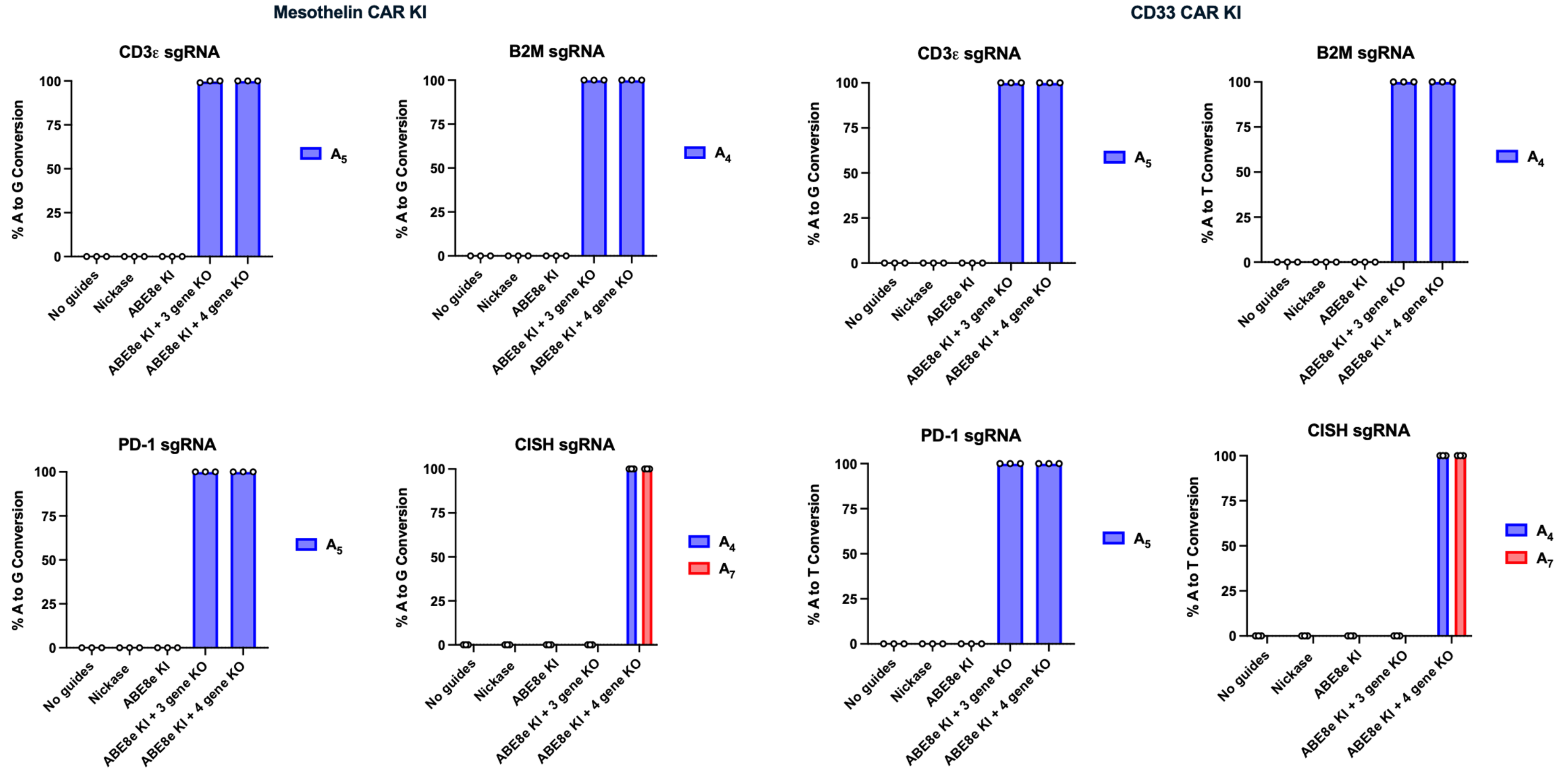


**Supplementary Figure 1.** Sanger sequencing analysis using EditR (https://moriaritylab.shinyapps.io/editr_v10/) of base editing outcomes at targeted loci. Simultaneous Mesothelin CAR integration with indicated degree of targeted KO **(Left, N=3 donors))** and CD33 CAR integration with indicated degree of targeted KO (**Right, N= 3 donors)**.

### Supplemental 2 - Flow Gating Strategy for surface detection of KO or CAR KI

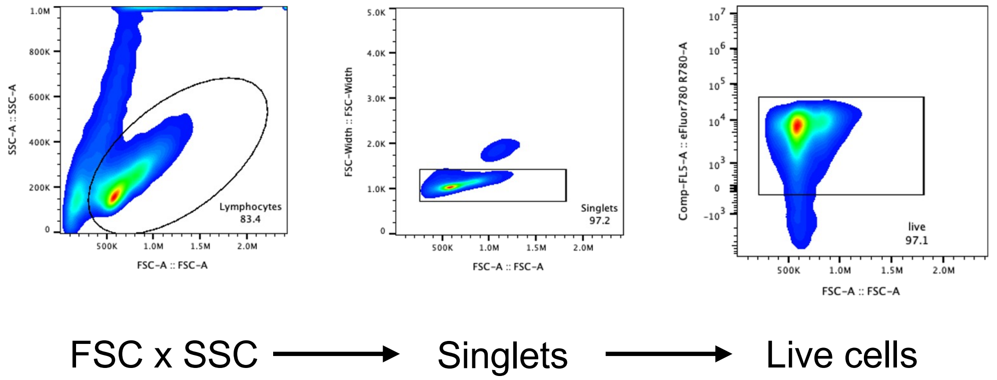


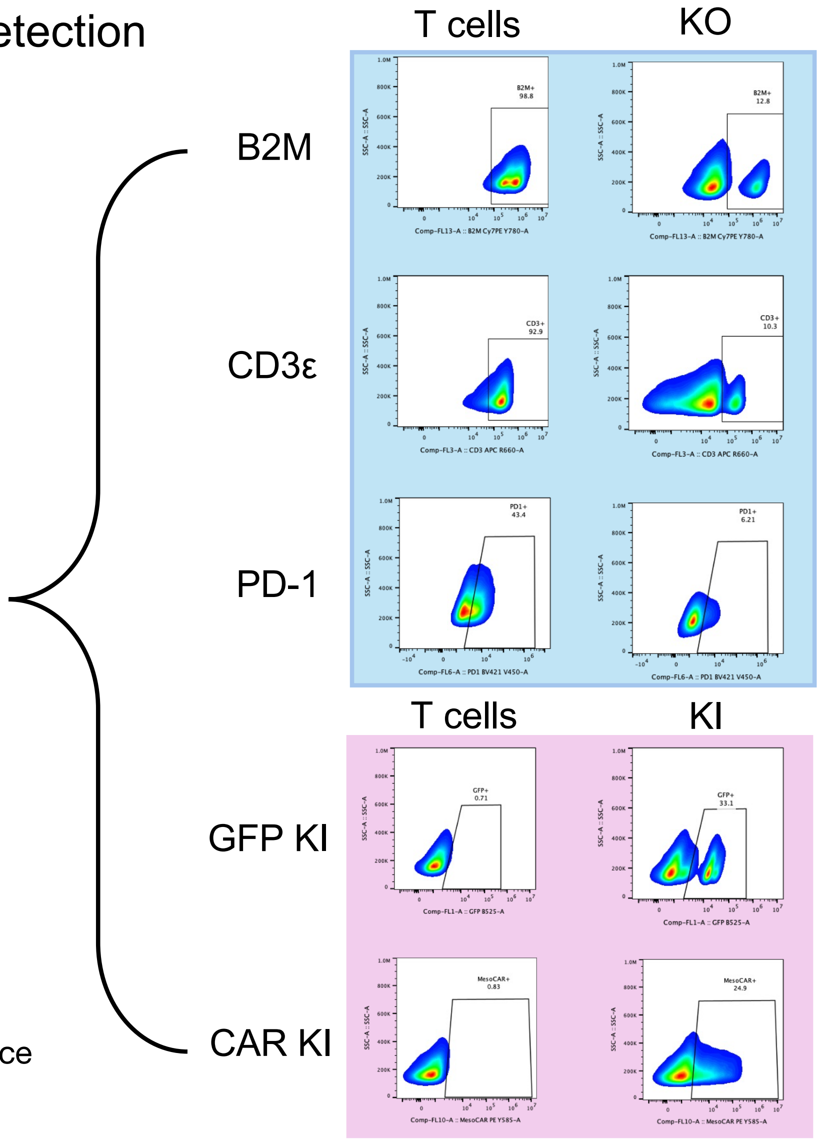


**Supplementary Figure 2.** Flow cytometry gating strategy to confirm surface protein knockout and/or CAR expression following INSERT engineering

### Supplemental Figure 3 - Target KO assessment via flow cytometry from engineering runs

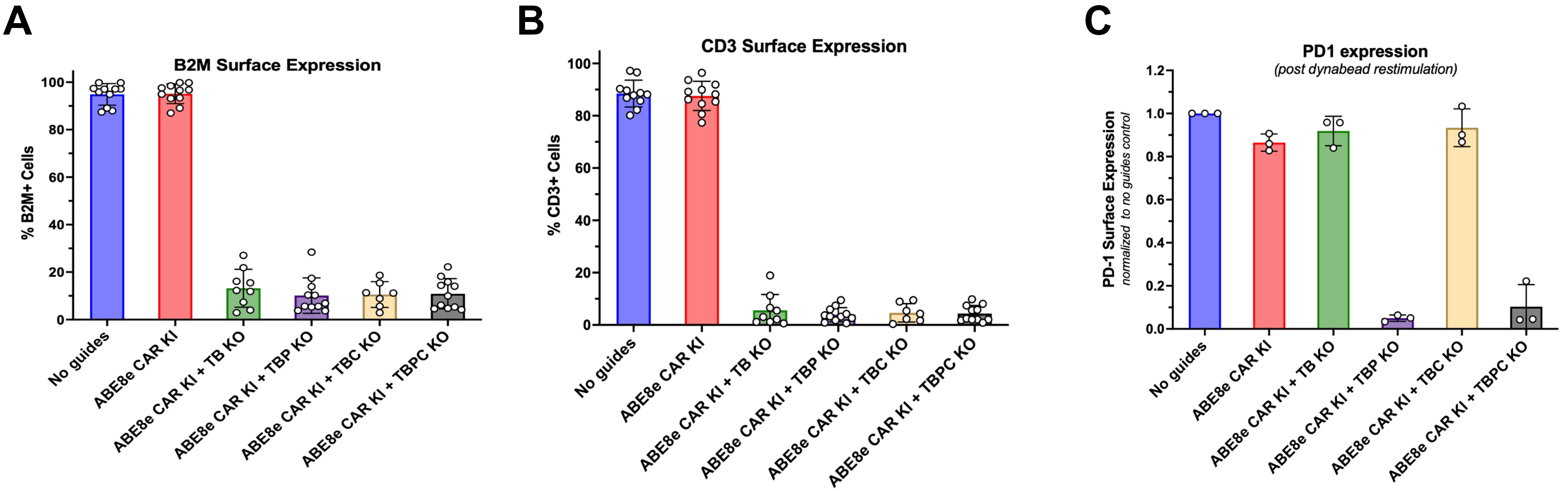


**Supplemental Figure 3:** Flow cytometry analysis of gene knockout efficiency in engineered T cells. (A) B2M surface expression, (B) CD3 surface expression, and (C) PD1 expression following dynabead restimulation across different experimental conditions. Cells were engineered with ABE8e base editor to knock-in CAR (CAR KI) alone or in combination with knockout of various targets:TB (CD3ε + B2M), TBP (CD3ε + B2M + PDCD1), TBC (CD3ε + B2M + CISH), or TBPC (CD3ε + B2M + PDCD1 + CISH). Expression levels were measured as percentage of positive cells (A,B) or normalized to no guide control (C). Each data point represents an individual donor, with bars showing mean values and error bars (+/-SD).

### Supplemental Figure 4 - CISH Western Blot and quantification of knockout

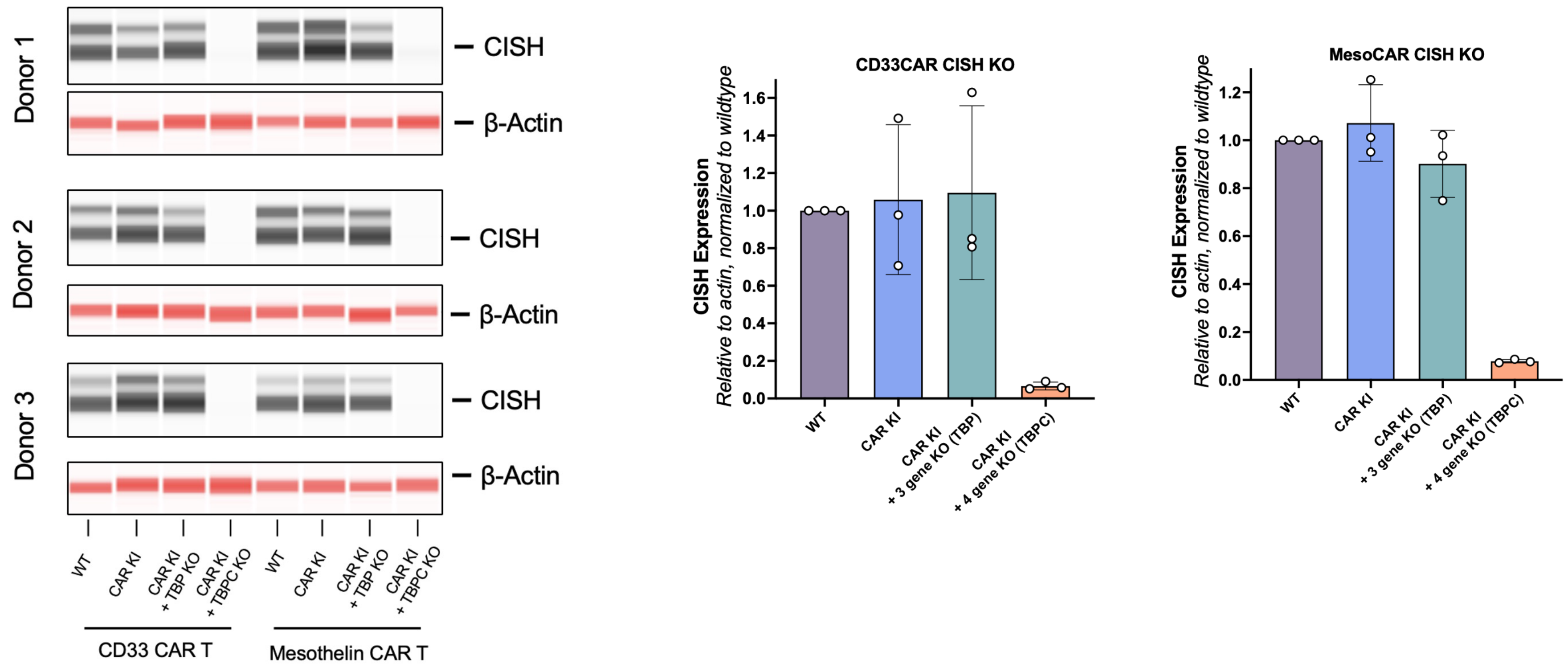


**Supplemental Figure 4: Western blot analysis of CISH protein expression in engineered T cells from three independent donors.** Left panels show representative western blots for CISH and β-Actin (loading control) across different experimental conditions: wild-type (WT), CAR knock-in (CAR KI), CAR KI with TBP knockout (CAR KI + TBP KO), and CAR KI with TBPC knockout (CAR KI + TBPC KO) in both CD33 CAR T cells and Mesothelin CAR T cells. Right panels show quantification of CISH expression normalized to β-Actin and relative to wild-type for CD33CAR CISH KO (Left) and MesoCAR CISH KO (Right). Each data point represents an individual sample, with bars showing mean values and error bars indicating standard deviation. TBP = CD3ε + B2M + PDCD1; TBPC = CD3ε + B2M + PDCD1 + CISH.

### Supplemental Figure 5 - T cell expansion data from INSERT T cells with increasing multiplex KO

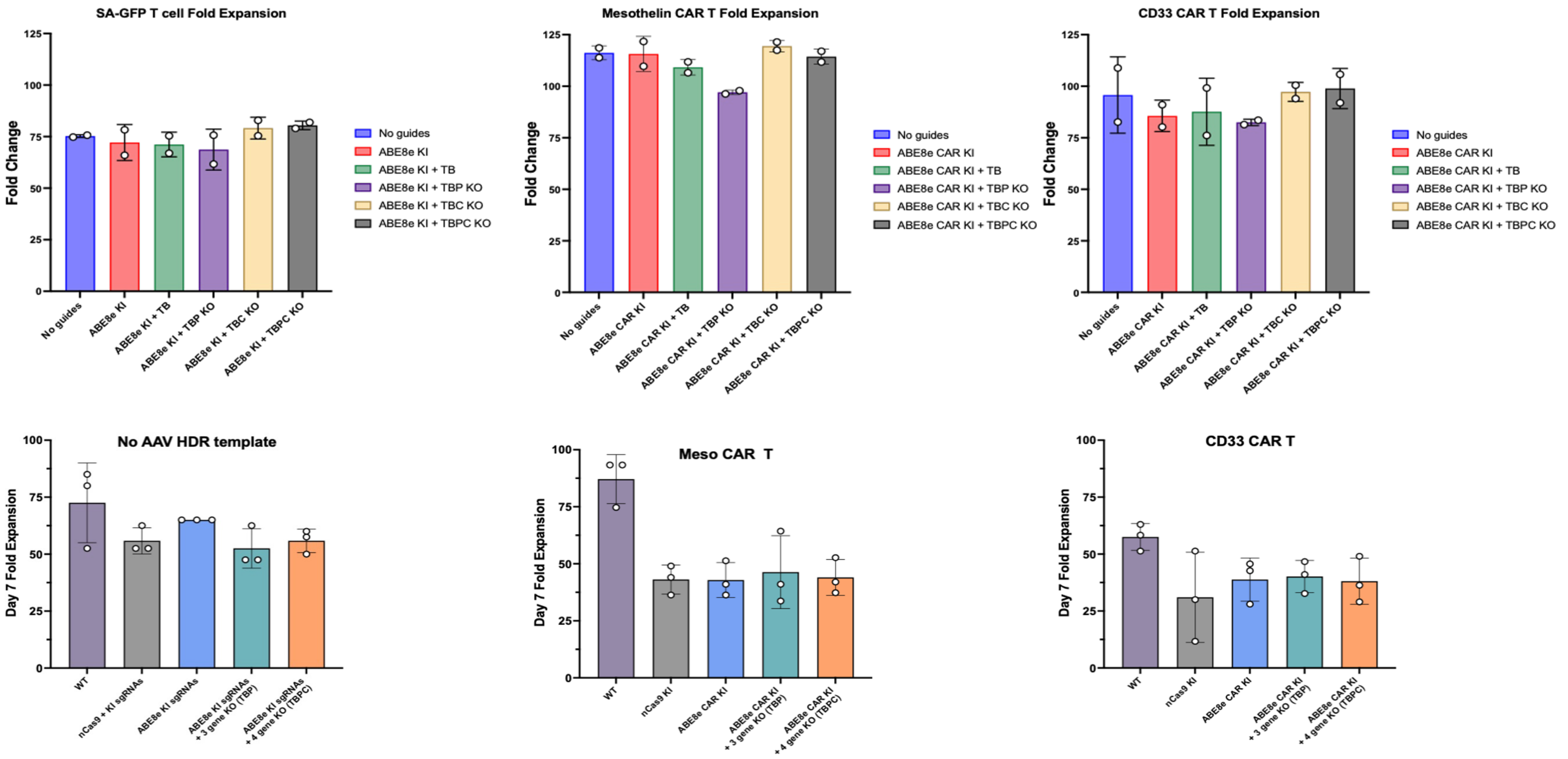


**Supplemental Figure 5: T cell expansion analysis of INSERT T cells with different multiplexed gene knockout combinations.** Upper panels show fold expansion of SA-GFP T cells (left), Mesothelin CAR T cells (middle), and CD33 CAR T cells (right) across various engineering conditions: No guides, ABE8e knock-in (ABE8e KI), ABE8e KI + TB (CD3ε + B2M), ABE8e KI + TBP KO (CD3ε + B2M + PDCD1), ABE8e KI + TBC KO (CD3ε + B2M + CISH), and ABE8e KI + TBPC KO (CD3ε + B2M + PDCD1 + CISH). Lower panels compare Day 7 fold expansion for different CAR T cell populations: No AAV HDR template (left), Mesothelin CAR T (middle), and CD33 CAR T (right) under engineering conditions. Each data point represents an individual sample, with bars showing mean values and error bars indicating standard deviation.

### Supplemental Table 1 - sgRNA sequences

| **Genome Target** | **Sequence** |
|---|---|
| *AAVS-1* #1 | CTTCCTAGTCTCCTGATATT |
| *AAVS-1* #2 | GTCACCAATCCTGTCCCTAG |
| *AAVS-1* #3 | CTTCCTGGTCTCCTGATATT |
| *AAVS-1* #4 | GTCGCCGGTCCTGTCCCTAG |
| *CD3e* (Ex.2 SD) | ACTCACCTGATAAGAGGCAG |
| *PDCD-1* (Ex.1 SD) | CACCTACCTAAGAACCATCC |
| *B2M* (Ex.1 SD) | CTTACCCCACTTAACTATCT |
| C*ISH* (Ex.2 SD) | CTCACCAGATTCCCGAAGGT |

Supplemental Table 1. sgRNA sequences used designed using. The table presents eight sgRNA sequences targeting different genomic regions: four targeting AAVS-1 (safe harbor locus) and four targeting genes for knockout: CD3ε (targeting exon 2 splice donor), PDCD-1 (targeting exon 1 splice donor), B2M (targeting exon 1 splice donor), and CISH (targeting exon 2 splice donor)

### Supplemental Table 2 - Sequencing primers

| Primer | Sequence |
|---|---|
| CD3e (Ex.2 SD) Forward | TAGTAAGTCTGCTGGCCTCC |
| CD3e (Ex.2 SD) Reverse | TGGTAAATGAGGCTCCTTGGT |
| PDCD-1 (Ex.1 SD) Forward | ATGCAGATCCCACAGGCG |
| PDCD-1 (Ex.1 SD) Reverse | CTCAGGGTAAGGGGCAGAG |
| B2M (Ex.1 SD) Forward | GGTGCCTGATATAGCTTGACACC |
| B2M (Ex.1 SD) Reverse | GACTCATTCAGGGTAGTATGGCC |
| CISH (Ex.2 SD) Forward | CCAGACAGAGAGTGAGCCAA |
| CISH (Ex.2 SD) Reverse | GGCAGAACTAAGCTTCTCCC |
| AAVS-1 #1 (5' AAVS1 KI site) Forward | CAGCTAGTCTTCTTCCTCCAAC |
| AAVS-1 #1 (5' AAVS1 KI site) Reverse | CTCCTGTTTCTCTGTTCCTGAC |
| AAVS-1 #2 (3' AAVS1 KI site) Forward | CCTGGTCATCACCCTGTATTG |
| AAVS-1 #2 (3' AAVS1 KI site) Reverse | CTCCATCGTAAGCAAACCTTAGA |

### Supplemental Table 3 - ddPCR primers and probes

| Primer or Probe Set | Sequence |
|---|---|
| RNAse P F Primer | AGA TTT GGA CCT GCG AGC G |
| RNAse P R Primer | GAG CGG CTG TCT CCA CAA GT |
| RNAse P Probe | /5HEX/TTC TGA CCT /ZEN/GAA GGC TCT GCG CG/3IABkFQ/ |
| AAVS-1 Junction F Primer | ATA TTC CCA GGG CCG GTT A |
| AAVS-1 Junction R Primer | GGG TGT GTC ACC AGA TAA GG |
| AAVS-1 5’ Junction Probe | /56-FAM/TG TGG CTC T/ZEN/G GTT CTG GGT ACT TT/3IABkFQ/ |
| AAVS-1 3’ Junction Probe | /56-FAM/CC CAC CTC C/ZEN/T GTT AGG CAG ATT /3IABkFQ/ |
| B2M 5' F Primer | CGT GTG AAC CAT GTG ACT TTG |
| B2M 3' R Primer | CAT ACA CAA CTT TCA GCA GCT TAC |
| CISH 5' F Primer | CCA GAC AGA GAG TGA GCC AA |
| CISH 3' R Primer | GGC AGA ACT AAG CTT CTC CC |
| CD3e 5' F Primer | GCA CTC ACT GGA GAG TTC TG |
| CD3e 3' R Primer | CCC TTT CCA CTC CAT CCT ACT |
| PD-1 5' F Primer | GGG CGG TGC TAC AAC TG |
| PD-1 3' R Primer | GGC TCT GGG ACA CCT GA |

### Supplemental Figure 6 - Flow diagram for ICS or surface CD107a analysis

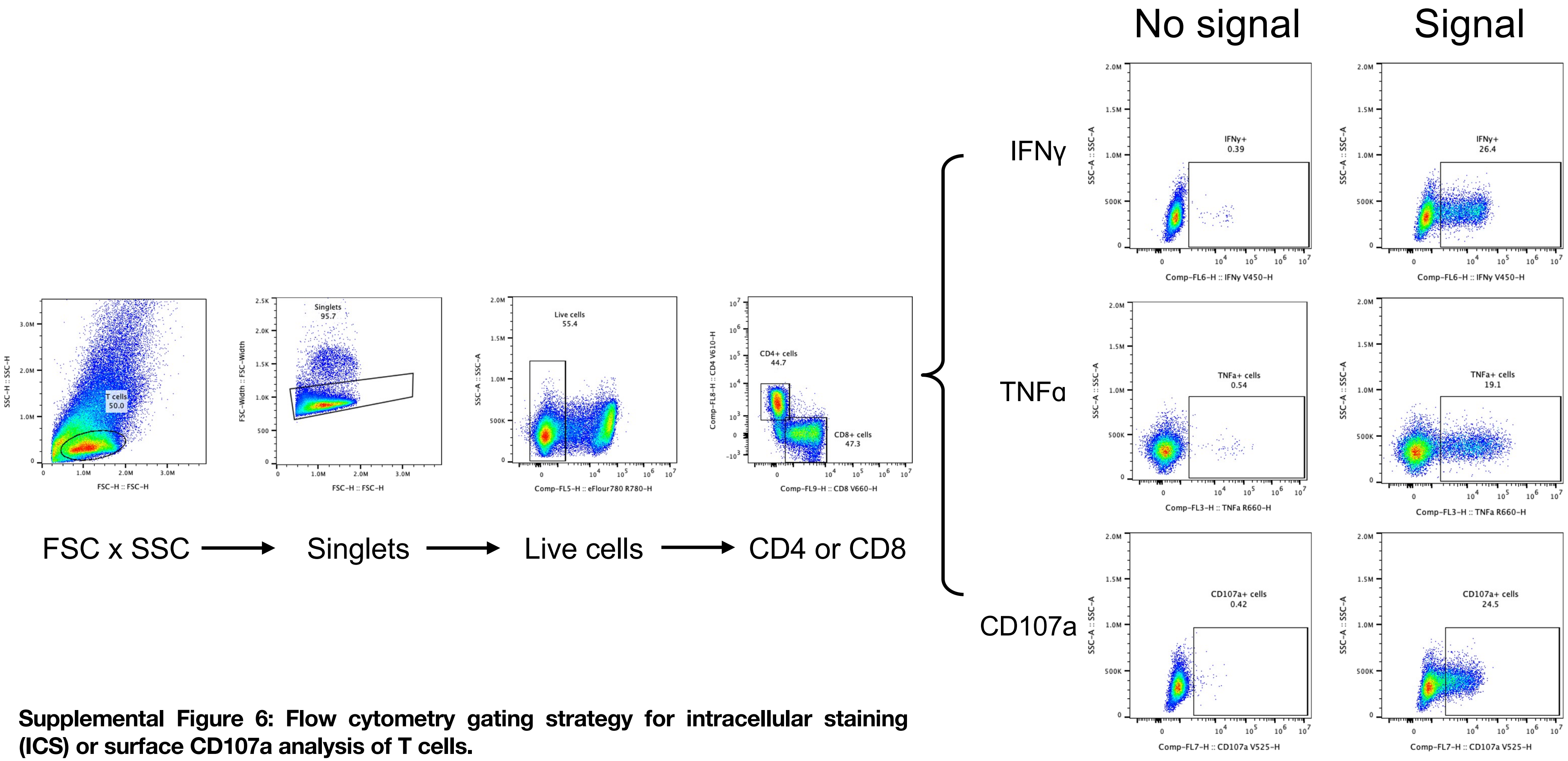


**Supplemental Figure 6: Flow cytometry gating strategy for intracellular staining (ICS) or surface CD107a analysis of T cells.**

### Supplemental Figure 7 - ICS results for CD33 CAR and GFP T

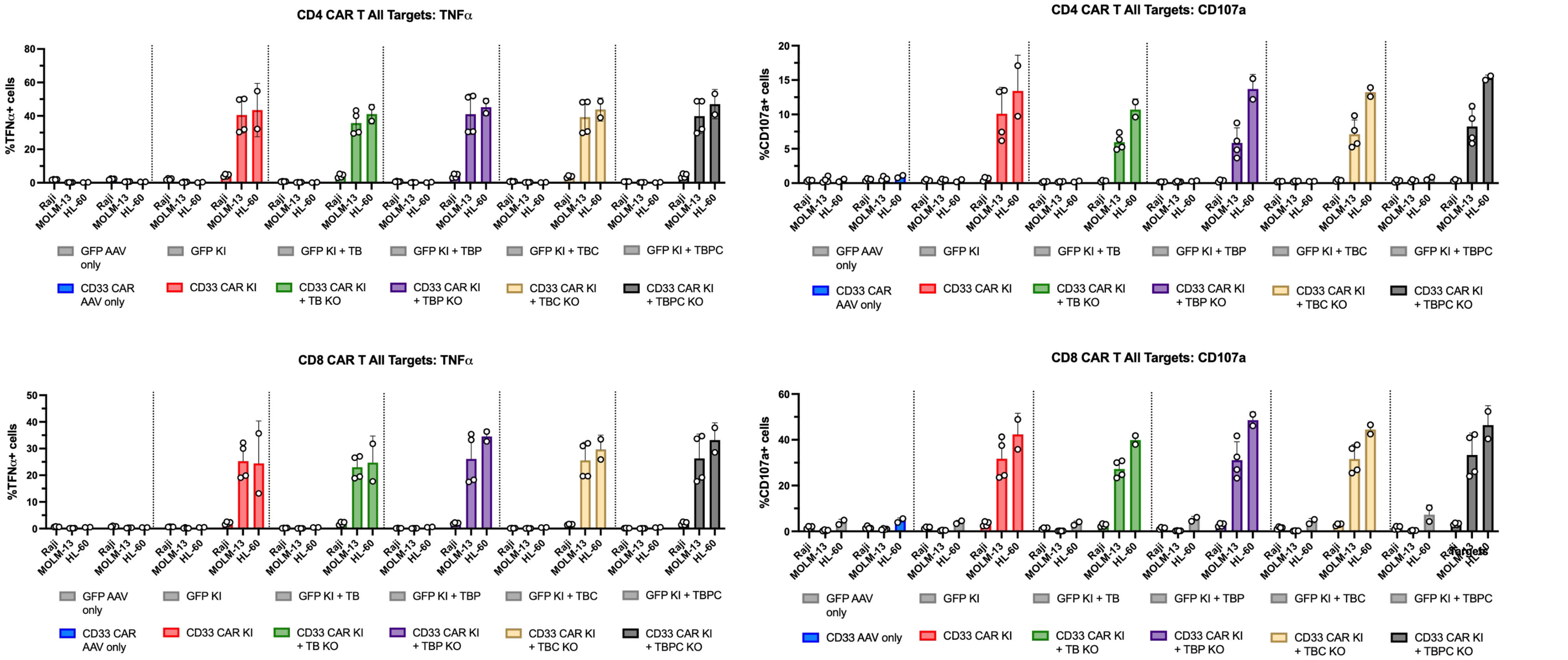


**Supplemental Figure 7:** Intracellular staining (ICS) analysis of CD33 CAR and GFP T cells following target cell stimulation. functional responses of CD4+ T cells (upper panels) and CD8+ T cells (lower panels) measured by TNFα production (left panels) and CD107a expression (right panels). Multiple engineering conditions: GFP AAV only, GFP KI, GFP KI + TB (CD3ε + B2M), GFP KI + TBP (CD3ε + B2M + PDCD1), GFP KI + TBC (CD3ε + B2M + CISH), GFP KI + TBPC (CD3ε + B2M + PDCD1 + CISH), CD33 CAR AAV only, CD33 CAR KI, CD33 CAR KI + TB KO, CD33 CAR KI + TBP KO, CD33 CAR KI + TBC KO, and CD33 CAR KI + TBPC KO. Cells were stimulated with MOLM13 cells expressing the target antigen (MOLM13-CD33) or control cells (MOLM13 HL-60). Each data point represents technical duplicated from N=2 donors, with bars showing mean values with SD.

### Supplemental Figure 8 - CD33 negative cell line killing assay

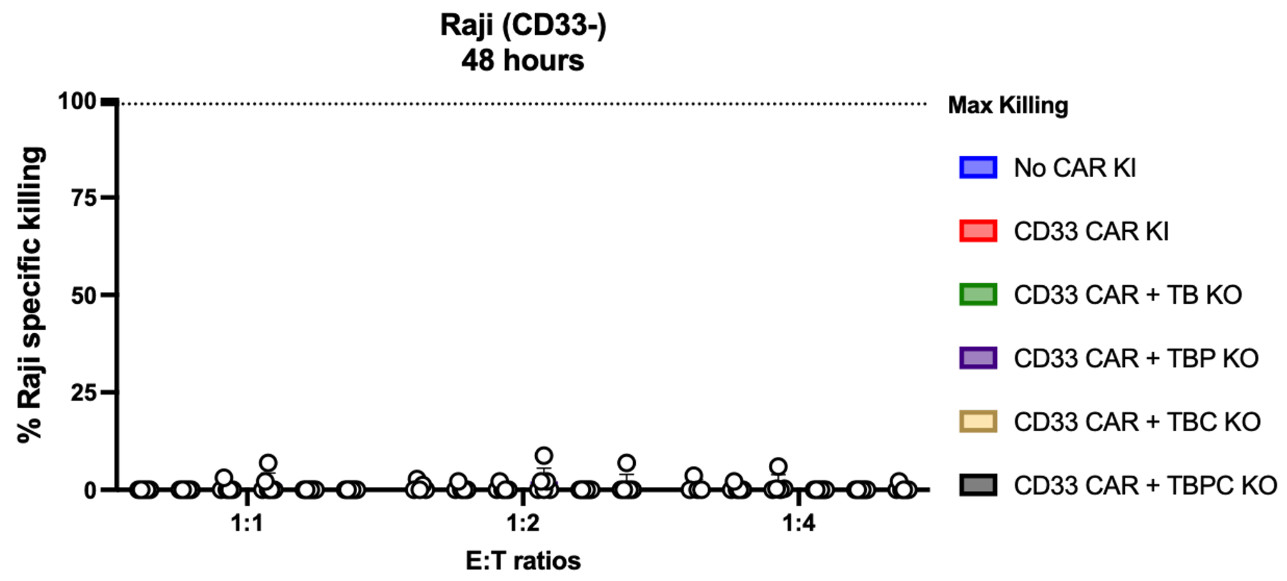


**Supplemental Figure 8:** Cytotoxicity assay using CD33-negative Raji cells as targets for engineered T cells. The graph shows percent specific killing of Raji (CD33-) cells after 48 hours of co-culture at different effector-to-target (E:T) ratios (1:1, 1:2, and 1:4). T cell populations: No CAR KI (blue), CD33 CAR KI (red), CD33 CAR + TB KO (CD3ε + B2M knockout, green), CD33 CAR + TBP KO (CD3ε + B2M + PDCD1 knockout, purple), CD33 CAR + TBC KO (CD3ε + B2M + CISH knockout, beige), and CD33 CAR + TBPC KO (CD3ε + B2M + PDCD1 + CISH knockout, grey).

Supplemental Figure 9 - Serial killing results from HL-60 targets

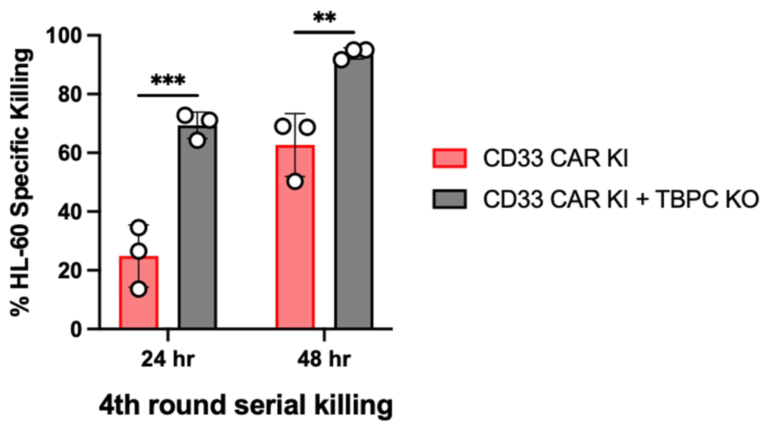


**Supplemental Figure 9.** Serial killing activity against HL-60 targets comparing CD33 CAR KI cells (orange bars) with CD33 CAR KI + TBPC KO cells (gray bars) at 24 and 48 hours during the 4th round of serial killing.

### Supplemental Figure 10 - MesoCAR cell killing assay

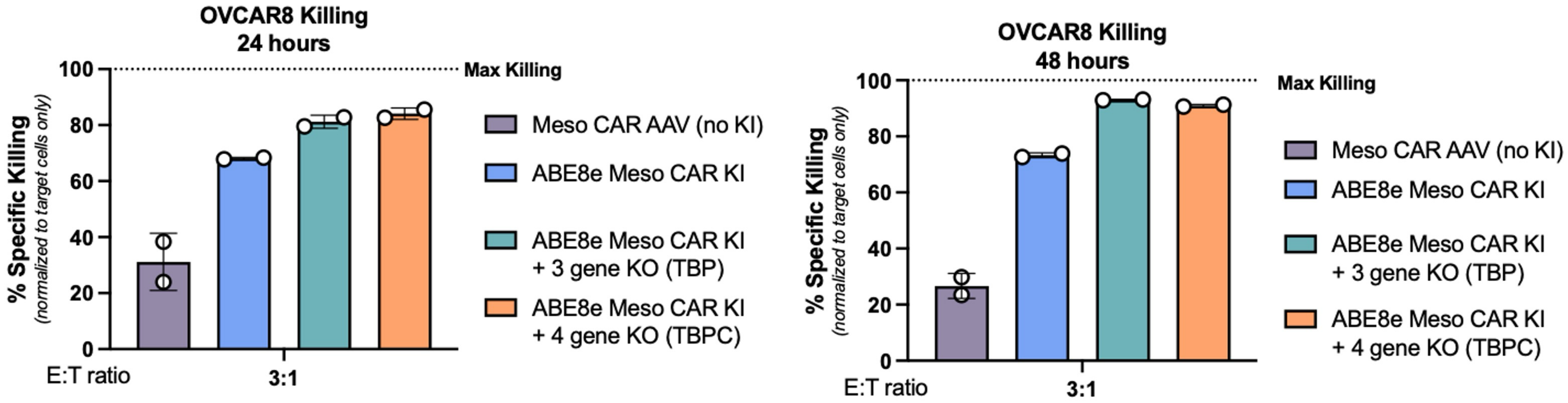


**Supplemental Figure 10.** OVCAR8 killing assay comparing cytotoxic activity of different MesoCAR cell types at 24 hours (left panel) and 48 hours (right panel) with an effector-to-target (E:T) ratio of 3:1. Four experimental conditions are shown: MesoCAR AAV (no KI) in purple, ABE8e MesoCAR KI in blue, ABE8e MesoCAR KI + 3 gene KO (TBP) (CD3ε + B2M + PDCD1 knockout, teal), and ABE8e MesoCAR KI + 4 gene KO (TBPC) (CD3ε + B2M + PDCD1 + CISH knockout, orange.)

Supplemental Figure 11 - metabolomics analysis of INSERT CD33 CAR T post first round killing

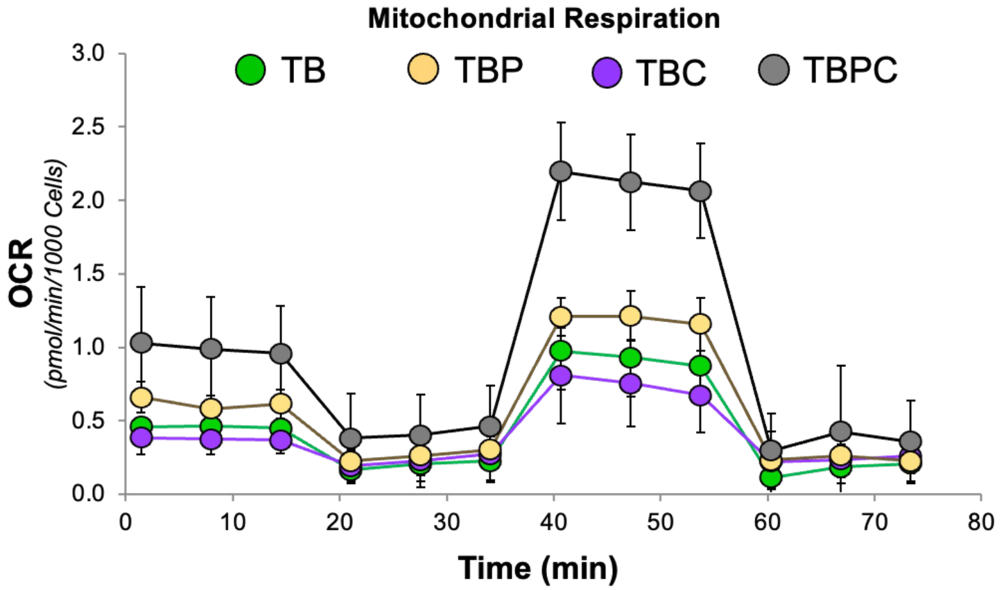


**Supplemental Figure 11. Metabolomics analysis of INSERT CD33 CAR T cells post first round killing, showing mitochondrial respiration profiles over time.** Displaying oxygen consumption rate (OCR, measured in pmol/min/1000 cells) as a function of time (minutes) for four different gene knockout combinations: T=CD3ε, B=B2M, P=PDCD1, and C=CISH.

### Supplemental Figure 12 - Sanger data and CISH western KO validation of INSERT CD19 CAR T

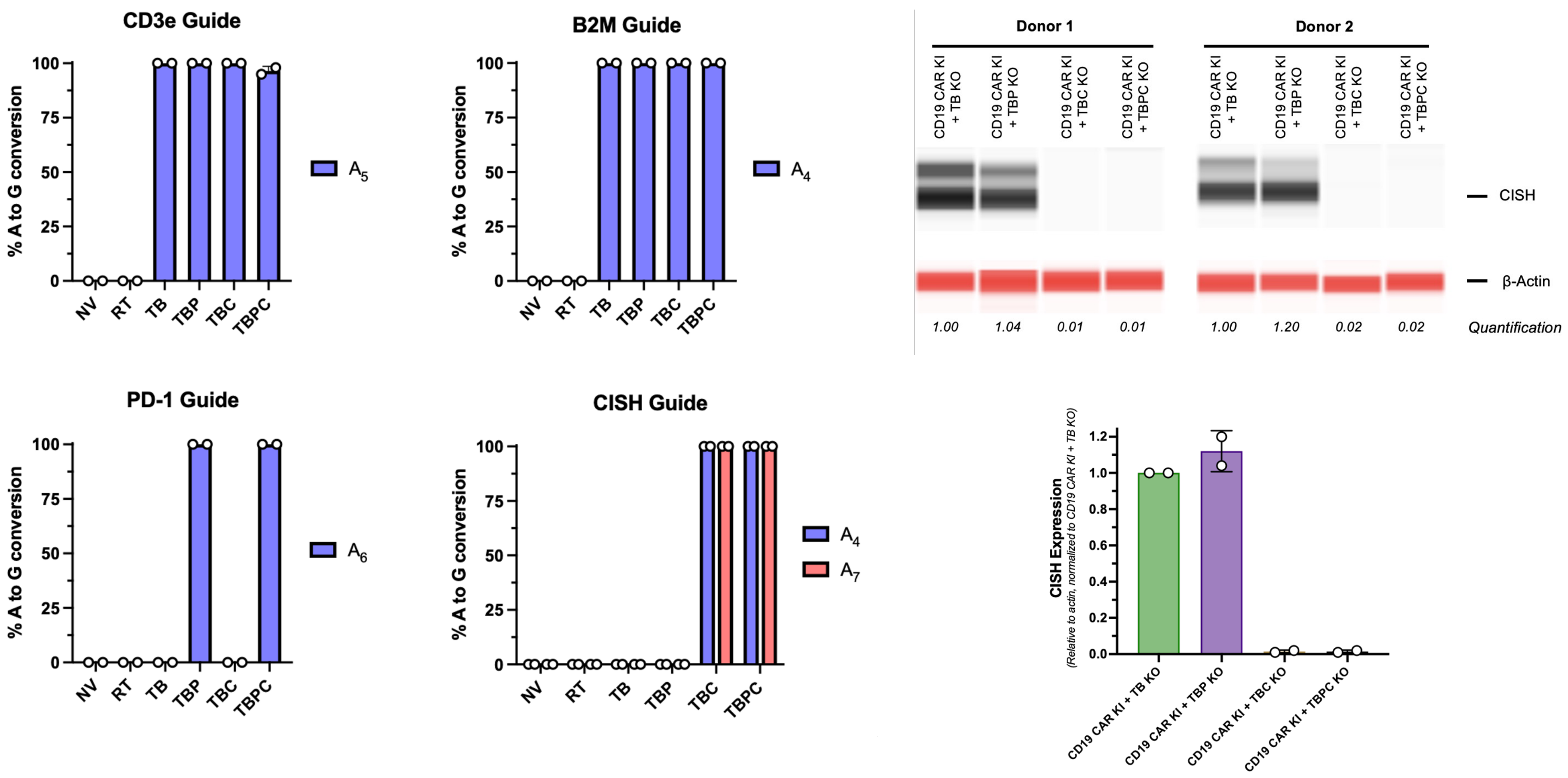

### Supplemental Figure 13 - INSERT CD19 CAR functional testing

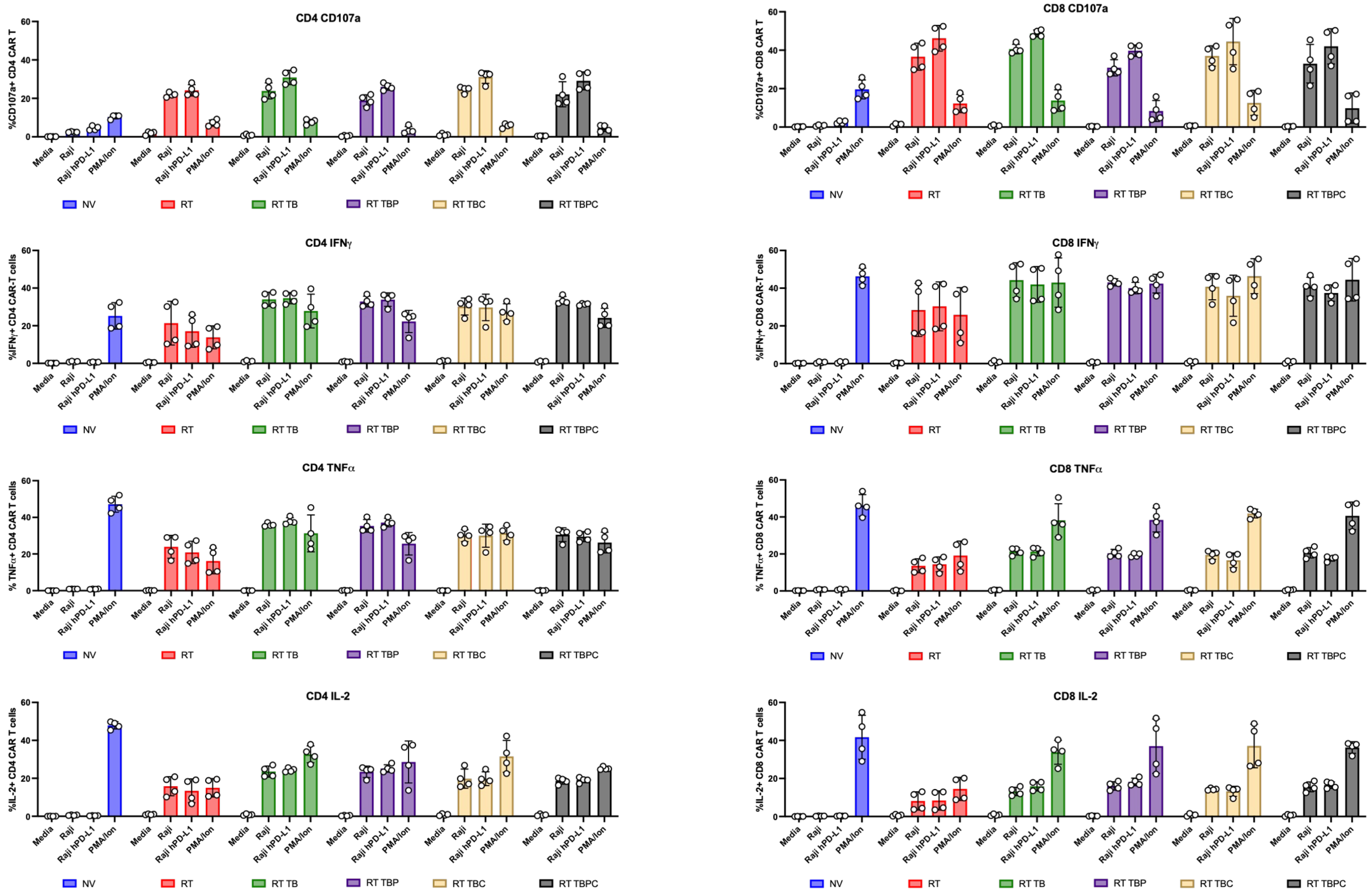

### Supplemental Figure 14 - CD19 CAR functional testing - Serial Killing

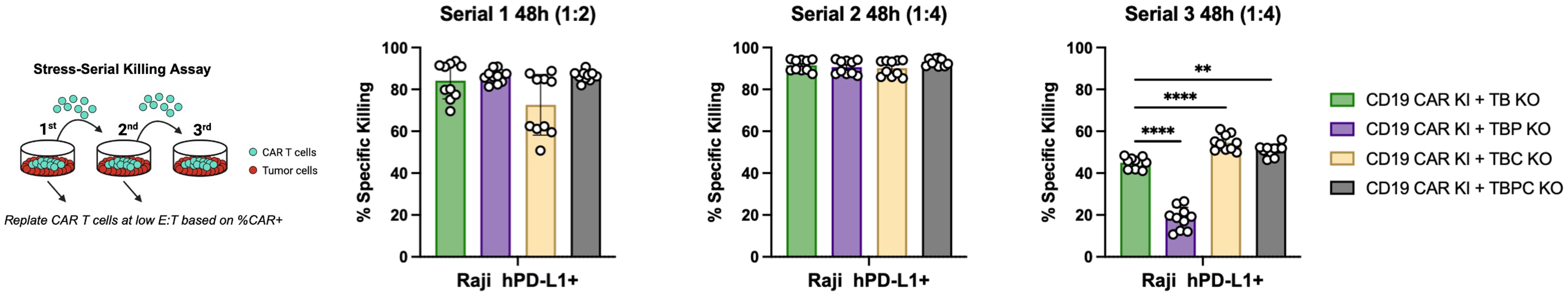


**Supplemental Figure 14. CD19 CAR functional testing through serial killing assays against Raji hPD-L1+ tumor cells.** The left panel illustrates the experimental design of the stress-serial killing assay where CAR T cells (teal) were replated with tumor cells (red) at low effector-to-target (E:T) ratios based on %CAR+ cells for successive rounds. The three right panels show specific killing percentages across three serial killing rounds (48 hours each) with different E:T ratios (1:2 for Serial 1, 1:4 for Serial 2 and 3). WIth Four different CD19 CAR KI conditions: TB KO (green), TBP KO (purple), TBC KO (beige), and TBPC KO (gray), where T=CD3ε, B=B2M, P=PDCD1, and C=CISH. ****p<0.0001, **p<0.01; One-way ANOVA followed by Tukey's multiple comparisons test against CD19 CAR KI + TB KO group. Replicates from N=2 donors shown.

Supplemental Figure 15 - Engineering circuit for nCas9

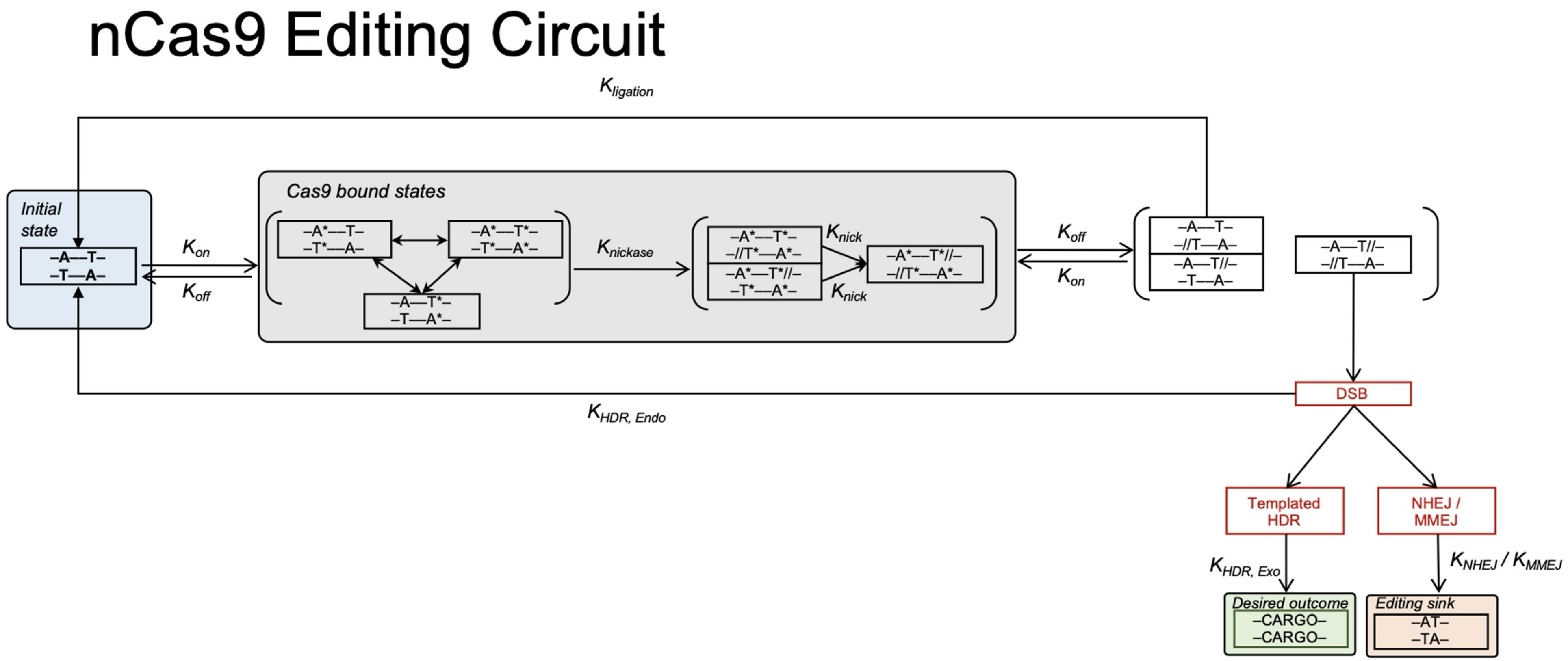


**Supplemental Figure 15. Working kinetic model of nCas9 DSB induction through staggered nicks**

#### Supplemental Figure 16 - Engineering circuit for ABE

### ABE Editing Circuit

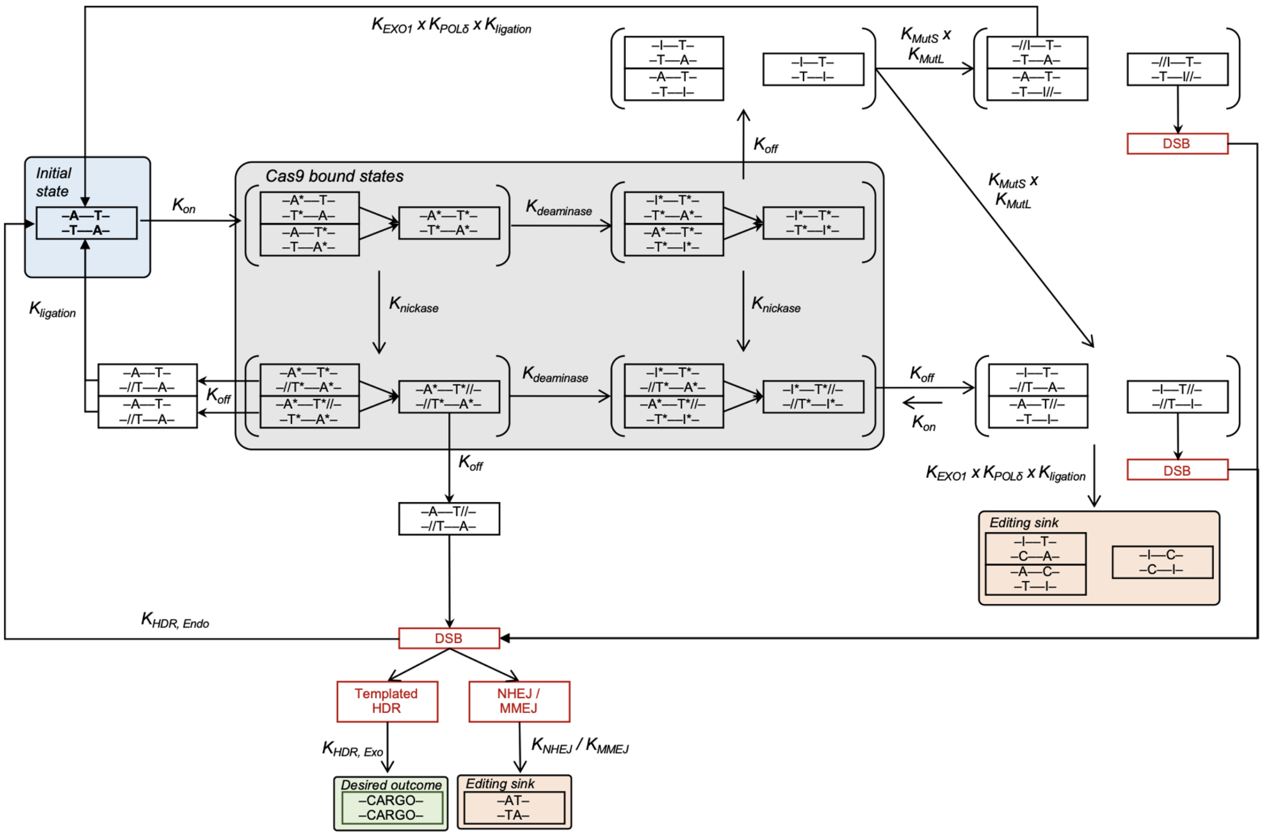


**Supplemental Figure 16. Working kinetic model of ABE DSB induction through staggered nicks**

### Supplemental Figure 17 - Engineering circuits for CBE

#### CBE Editing Circuit

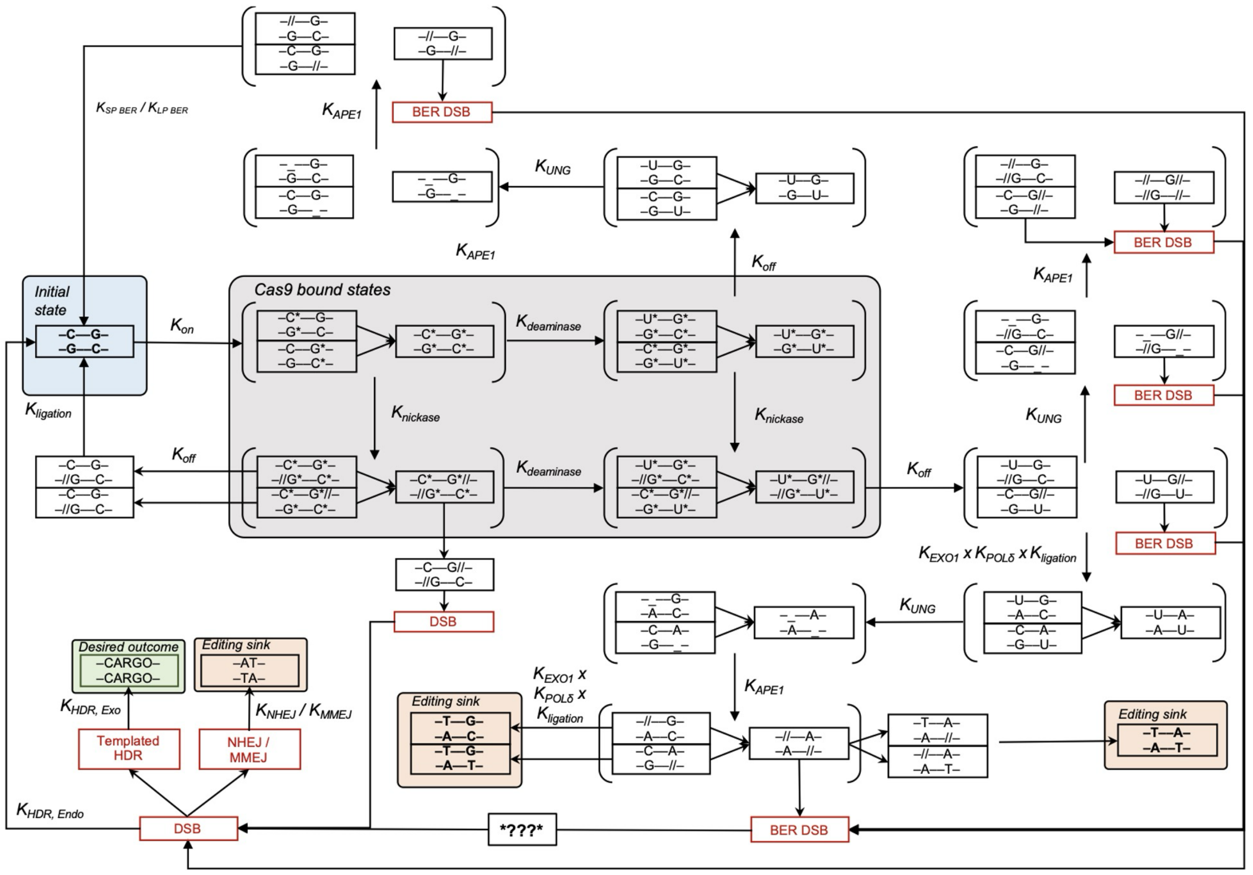


**Supplemental Figure 17. Working kinetic model of CBE DSB induction through staggered nicks**
